# Evolutionary history and functional divergence of hydroxycarboxylic acid receptors in primates

**DOI:** 10.64898/2026.01.23.701403

**Authors:** Juan C. Opazo, L. Felipe Barros, Kattina Zavala, Rodrigo Maldonado, Gonzalo A. Mardones

## Abstract

Hydroxycarboxylic acid receptors (HCARs) are class A G-protein-coupled receptors that function as metabolic sensors. This receptor family includes three members (HCAR1, HCAR2, and HCAR3) expressed in metabolically active tissues and immune cells, where they link cellular metabolic status to physiological responses. This study aims to elucidate the evolutionary history of the most recently originated members of the HCAR gene family, namely HCAR2 and HCAR3, in primates. According to our phylogenetic analyses, the duplicative history of these genes involved multiple independent duplication events during ape evolution. Thus, most ape lineages possess independently originated duplicated copies, while non-ape primates retain the ancestral condition of a single-copy gene (HCAR2/3). Our analyses further indicate that this single-copy gene in non-ape primates is functionally equivalent to HCAR2, suggesting that the primary functional innovation in apes is associated with the physiological roles of HCAR3. Finally, gene expression analyses reveal that major divergence in tissue expression occurred after the initial duplication event that generated HCAR1 and the HCAR2/3 lineage, whereas HCAR2 and HCAR3 exhibit substantial overlap in their expression profiles. Thus, the more refined and context-dependent regulation of lipid metabolism that provides the HCAR3 receptor seems to have originated multiple times during the evolutionary history of apes.

## Introduction

The study of gene family evolution is rapidly advancing field that has benefited greatly from increasing availability of freely accessible genomic data. In primates, the growing number of high-quality whole-genome sequences (Kuderna et al. 2023; Upham and Landis 2023; Dyer et al. 2025; Sayers et al. 2025) together with well-resolved phylogenetic relationships (Perelman et al. 2011; Shao et al. 2023; S. Wang et al. 2025) provides an ideal framework for investigating the evolutionary dynamics of gene families. This is particularly important given that the evolutionary histories of a substantial proportion of genes remain poorly characterized (Altenhoff et al. 2016). Accurately defining homologous relationships is therefore essential, not only for understanding how genes are related to one another, but also because such definitions have important downstream consequences when comparing genes across species, where analyses often rely on the assumption that 1:1 orthologous genes or proteins are being compared.

Hydroxycarboxylic acid receptors (HCARs) are class A G-protein-coupled receptors that detect metabolic intermediates and primarily signal through Gi/o proteins (Offermanns 2014; Offermanns 2017). In humans, the family includes HCAR1, HCAR2, and HCAR3. HCAR1 is activated by lactate and is expressed in adipocytes, muscle, and other metabolically active tissues, where it inhibits lipolysis and modulates metabolic adaptation under conditions of high glycolytic flux (Liu et al. 2009; Offermanns 2014; Offermanns 2017). HCAR2 is activated by β-hydroxybutyrate, niacin, and short-chain fatty acids, regulating lipolysis, energy homeostasis, and anti-inflammatory responses in metabolic and immune tissues (Taggart et al. 2005; Offermanns 2014; Offermanns 2017). Finally, HCAR3 responds to 3-hydroxyoctanoic acid and metabolites derived from lactic acid bacteria, such as D-phenyllactic acid, thereby linking lipid metabolism and dietary fermentation products to host immune and metabolic regulation (Ahmed et al. 2009; Offermanns 2014; Offermanns 2017; Peters et al. 2019). Structural analyses indicate that ligand binding in HCAR2 and HCAR3 occurs within a conserved orthosteric pocket, where polar interactions anchor the ligand carboxyl group and hydrophobic contacts stabilize ligand binding (Park et al. 2023; Suzuki et al. 2023; Ye et al. 2024; J. Wang et al. 2025). While the overall pocket architecture is highly similar between both receptors, subtle differences in pocket composition modulate ligand selectivity and binding stability (Park et al. 2023; Suzuki et al. 2023; Ye et al. 2024; J. Wang et al. 2025).

From an evolutionary perspective, although it is well recognized that HCARs form a gene family, their duplicative history is not well resolved. In a relatively recent study, Peters et al. (2019) reconstructed the evolutionary history of HCARs and proposed that HCAR2 and HCAR3 originated, through a gene duplication event, in the last common ancestor of apes, dated to approximately 28.8–19.5 million years ago. At the same time, they suggested that non-ape primates possess an HCAR2 gene in their genomes. This interpretation is problematic, as it is inconsistent with the proposed timing of the duplication event that gave rise to HCAR2 and HCAR3 in the ape ancestor (Peters et al. 2019).

This study aims to further elucidate the evolutionary history of the most recently originated members of the HCAR gene family, namely HCAR2 and HCAR3. Our results indicate that the duplication history of these genes is more dynamic than previously appreciated, with multiple, independent duplication events occurring during ape evolution. As a consequence, most ape lineages harbor independently derived duplicated copies, whereas non-ape primates retain the ancestral single-copy condition (HCAR2/3). Structural and sequence analyses suggest that the single-copy gene present in non-ape primates is functionally equivalent to HCAR2, implying that the primary functional innovation following duplication is associated with HCAR3. Finally, gene expression analyses reveal that major divergence in tissue expression occurred after the earlier duplication that gave rise to HCAR1 and the HCAR2/3 lineage, whereas HCAR2 and HCAR3 themselves exhibit substantial overlap in their expression profiles.

## Materials and Methods

### DNA Sequences

We manually annotated the hydroxycarboxylic acid receptor 2 (HCAR2, Q8TDS4) and hydroxycarboxylic acid receptor 3 (HCAR3, P49019) genes in representative species from all major primate groups (Supplementary Table S1). Genomic fragments containing HCAR genes were retrieved from the National Center for Biotechnology Information (NCBI) database (Sayers et al. 2025), based on conserved synteny. Each fragment included the HCAR genes along with its flanking genes: the 5′ flanking gene, Density-regulated protein (DENR, O43583), and the 3′ flanking gene, Kinetochore-associated protein 1 (KNTC1, P50748). We then manually curated the existing gene annotations or performed de novo annotation by aligning known exon sequences from closely related species to the target genomic fragments. These comparisons were carried out using the Blast2Seq v2.5 program with default parameters (Tatusova and Madden 1999).

Nucleotide sequences, including 500 bp upstream, the full coding region, and 500 bp downstream, were aligned using MAFFT v7 (Katoh and Standley 2013), using the L-INS-i strategy selected automatically by the software. To identify the best-fitting model of molecular evolution, we applied the model selection routine in IQ-TREE v1.6.12 (Kalyaanamoorthy et al. 2017), which selected TIM3e+R3 as the best model. Phylogenetic inference was conducted using a maximum likelihood approach in IQ-TREE v1.6.12 (Trifinopoulos et al. 2016). We performed 20 independent tree searches, varying the perturbation strength parameter (-pers: 0.3, 0.5, 0.7, and 0.9) and setting the number of unsuccessful iterations to stop parameter (-nstop) to 1000 (default: 100). The tree with the highest log-likelihood score (−12576.162; -pers 0.3; -nstop 1000) was selected. Node support was evaluated using the aBayes test (Anisimova et al. 2011) and the ultrafast bootstrap method with 1,000 replicates (Hoang et al. 2018).

### Assessment of Conserved Synteny

We examined genes found up- and downstream of hydroxycarboxylic acid receptor genes in species representative of primates. Synteny assessments were conducted using the NCBI database (Sayers et al. 2025). Our analyses included the following species: human (*Homo sapiens*), Western lowland gorilla (Gorilla gorilla gorilla), Sumatran orangutan (*Pongo abelii*), siamang (*Symphalangus syndactylus*), Eastern hoolok gibbon (Hoolock leuconedys), northern white-cheeked gibbon (*Nomascus leucogenys*), crab-eating macaque (*Macaca fascicularis*), *Cercopithecus sabaeus*, Golden snub-nosed monkey (*Rhinopithecus roxellana*), white-tufted-ear marmoset (*Callithrix jacchus*), white-faced saki (*Pithecia pithecia*), Philippine tarsier (*Carlito syrichta*), coquereĺs sifaka (*Propithecus coquereli*), Ring-tailed lemur (*Lemur catta*) and slow loris (*Nycticebus coucang*).

### Structural Analyses

Protein structure modeling of non-ape primates, full-length HCAR2/3 receptors without bound ligands, was performed using the AlphaFold3 server (https://alphafoldserver.com) (Abramson et al. 2024). Structural models of non-ape primate HCAR2/3 receptors bound to ligands were generated using Protenix, an advanced AI-based, open-source reproduction of AlphaFold3 (https://protenix-server.com/model/prediction) (Chen et al. 2025). Ligand-protein interaction diagrams were generated using LigPlot^+^ v2.2 (Laskowski and Swindells 2011). Root-mean-square deviation (RMSD) values and structural figures were prepared using the PyMOL Molecular Graphics System, v3.0.4 (Schrödinger, LLC). For consistency, RMSD values were calculated using amino acids 8-302 when comparing human HCAR2 and HCAR2/3 models, and amino acids 17-300 when comparing HCAR3 and HCAR2/3 models. Statistical significance between mean RMSD values was assessed using a two-tailed, paired *t*-test, with a *P*-value < 0.05 considered statistically significant (*P* = 0.0003).

## Results and Discussion

### Independent Evolutionary Origins of HCAR2 and HCAR3 Gene Lineages in Apes

We conducted a phylogenetic reconstruction to investigate the evolutionary history of the hydroxycarboxylic acid receptor 2 (HCAR2) and 3 (HCAR3) genes in primates. Our tree topology recovered the sister group relationships among the main primate groups, consistent with the most recent evolutionary hypotheses (Perelman et al. 2011; Shao et al. 2023),(Perelman et al. 2011; Shao et al. 2023),(Perelman et al. 2011; Shao et al. 2023). Our analyses revealed several gene expansion events in apes, leading to the emergence of independent gene duplicates (Fig. 1). In contrast, representative species of strepsirhines, tarsiers, New World monkeys, and Old World monkeys retain the ancestral condition of a single-copy gene (HCAR2/3) (Fig. 1). Accordingly, the last common ancestor of primates, dated to approximately 74 million years ago (Kumar et al. 2017), possessed a repertoire of a single-copy gene (HCAR2/3).

**Figure 1.**
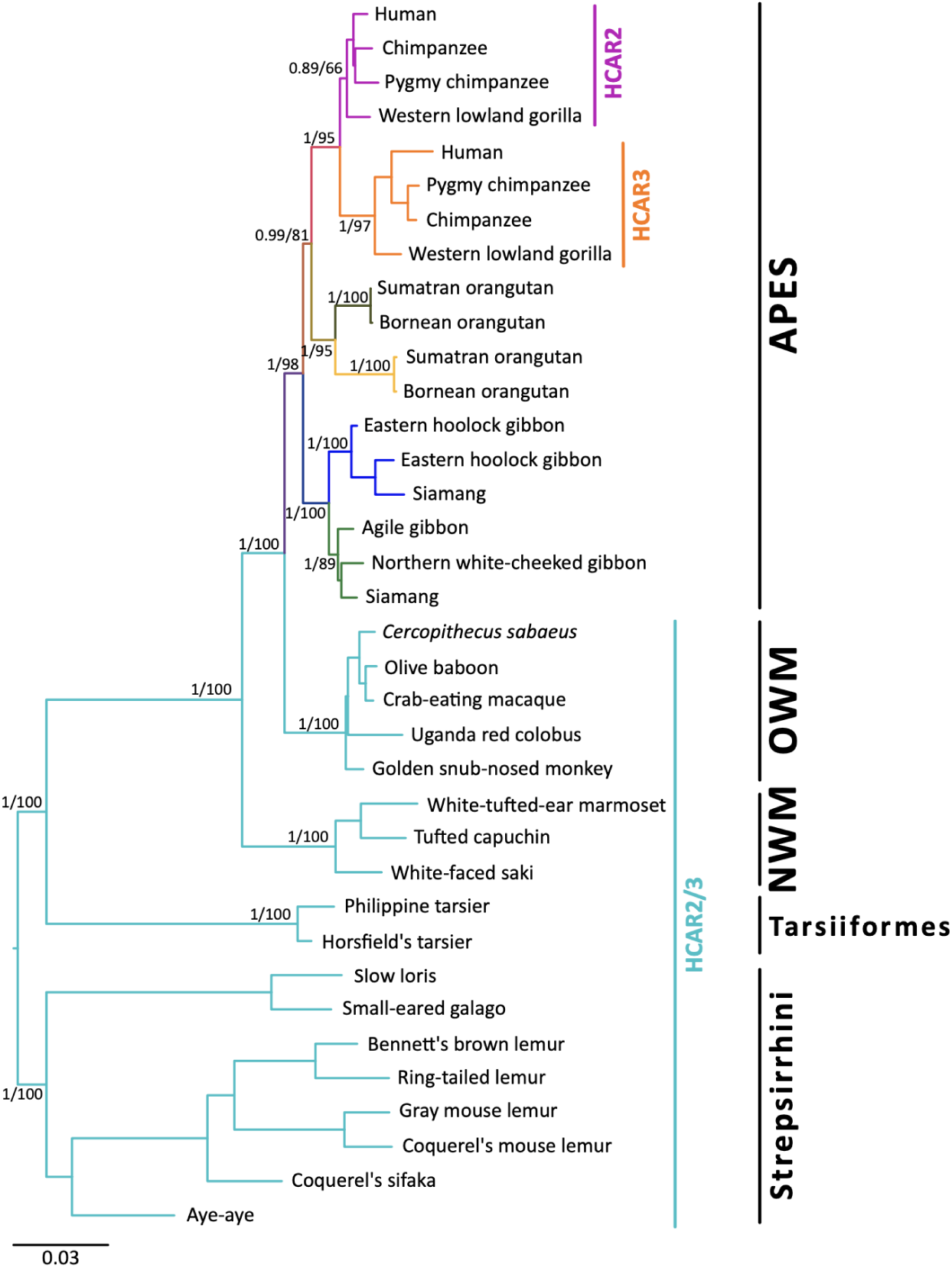
Phylogenetic relationships among hydroxycarboxylic acid receptor 2 and 3 (HCAR2 and HCAR3) genes in primates. Numbers on the nodes correspond to support from the aBayes and ultrafast bootstrap values. The scale denotes substitutions per site, and colors represent gene lineages.

In apes, the HCAR2/3 gene underwent independent duplication events in gibbons, orangutans and in the common ancestor of humans, chimpanzees, pygmy chimpanzees and gorillas (Fig. 1). In humans, the HCAR2 gene corresponds to the copy located on the 5′ side of the kinetochore-associated protein 1 (KNTC1) flanking gene, whereas HCAR3 is situated in the middle of the HCAR gene cluster, on the 5′ side of HCAR1. This syntenic organization is conserved across primates. The case of lesser apes (gibbons) is particularly notable due to lineage-specific patterns of gene retention and loss. For example, the siamang (*Symphalangus syndactylus*) has retained both gene copies derived from an ancestral duplication event (Fig. 1). In contrast, other gibbon species, such as the agile gibbon (*Hylobates agilis*) and the northern white-cheeked gibbon (*Nomascus leucogenys*), have retained only a single-gene copy (Fig. 1). Interestingly, the eastern hoolock gibbon (*Hoolock leuconedys*) possesses two gene copies, but both belong to the same gene lineage, suggesting a more recent duplication event within that lineage (Fig. 1). Overall, our phylogenetic analyses do not support the previous phylogenetic reconstruction reported by Peters et al. (2019), which propose a single duplication event in the ape ancestor, dated to approximately 28.8–19.5 million years ago, to explain the origin of HCAR2 and HCAR3. In contrast, our phylogeny suggests a more complex duplicative history, requiring multiple duplication events to account for the observed gene tree (Fig. 1). Moreover, Peters et al. (2019) suggested that non-ape primates, e.g., Old World and New World monkeys, possess an HCAR2 gene in their genomes, however, this inference is not supported by their results.

Thus, according to our gene tree/species tree reconciliation analyses, the HCAR homologous relationships are not straightforward. Although most ape species possess a repertoire of two genes, they are not always 1:1 orthologous. HCAR2 genes are 1:1 orthologous in the case of humans, chimpanzees, and gorillas; the same applies to HCAR3. However, HCAR2 and HCAR3 of this group of species are not 1:1 orthologous in the other ape species, as they originated from independent gene duplication events. This distinction is important to consider when making comparative analyses. It becomes especially relevant when comparing functional data with non-ape species (e.g., macaques, marmosets), which retain the ancestral condition of a single-copy gene. The caution required when comparing gene repertoires that originated through independent duplication events is not unusual in nature and fundamentally reflects the way in which evolution operates. Rather than being seen solely as a difficulty, such cases offer valuable opportunities to explore the evolutionary consequences of convergent events. The literature provides several notable examples of gene families that have expanded independently in different groups. For instance, the β-globin gene clusters originated independently in mammals, birds, crocodiles, turtles, and squamates (Goodman et al. 1987; Hoffmann et al. 2018), as well as separately in monotremes and therian mammals (Opazo et al. 2008). Similarly, the growth hormone gene family arose independently in New World monkeys and catarrhine primates (Li et al. 2005). Independent origins have also been documented for selective ion channels across distinct groups of gnathostomes (Flores-Aldama et al. 2020) and for lymphocyte receptors in gnathostomes and cyclostomes (Huang et al. 2024). These examples underscore the importance of first reconstructing the evolutionary history of gene families before attempting to compare their biological properties across taxonomic groups. Without this context, interpretations may be misleading, especially considering that the evolutionary trajectories of the vast majority of gene families remain unresolved (Altenhoff et al. 2016).

### Molecular Signatures of HCAR2 and HCAR3 Paralogues

The HCAR2 and HCAR3 genes, found in humans, chimpanzees, pygmy chimpanzees, and gorillas, originated from a gene duplication event in their common ancestor approximately 8.6 million years ago (Kumar et al. 2017). Due to this relatively recent divergence, their amino acid sequences remain highly similar, with pairwise identity values ranging from 95.03% (human HCAR3 vs. gorilla HCAR2) to 100% (bonobo HCAR2 vs. chimpanzee HCAR2).

Human HCAR2 and HCAR3 are highly structurally similar seven-transmembrane helical bundle G-protein coupled receptors (Ye et al. 2024). Although the human HCAR2 and HCAR3 proteins differ at 18 amino acid positions, a multispecies alignment reveals that 12 of these residues consistently and unambiguously distinguish the two receptors, differences that likely contribute to their distinct ligand-binding specificities (Fig. 2). Experimental and structural data have shown that the HCAR2 binding pocket is polar and compact, whereas the HCAR3 pocket is more hydrophobic and spacious (Park et al. 2023; Liu et al. 2024; Ye et al. 2024). These structural differences, arising from specific amino acid substitutions within the ligand-binding region and adjacent loops, confer unique ligand selectivity on each receptor. Among the distinguishing residues, six are located within transmembrane (TM) domains: one in TM1 (position 27), two in TM2 (positions 83 and 86), two in TM3 (positions 103 and 107), and one in TM4 (position 142) (Figs. 2 and 3). The remaining distinguishing residues lie between transmembrane domains, specifically at positions 91, 167, 168, 173, 176, and 178, with most situated on the extracellular side (Figs. 2 and 3). Notably, five of the distinguishing residues (positions 27, 142, 167, 168, and 173) are located at sites distant from the ligand-binding pocket (Fig. 3), suggesting that these residues may contribute to additional functional differences between the paralogues.

**Figure 2.**
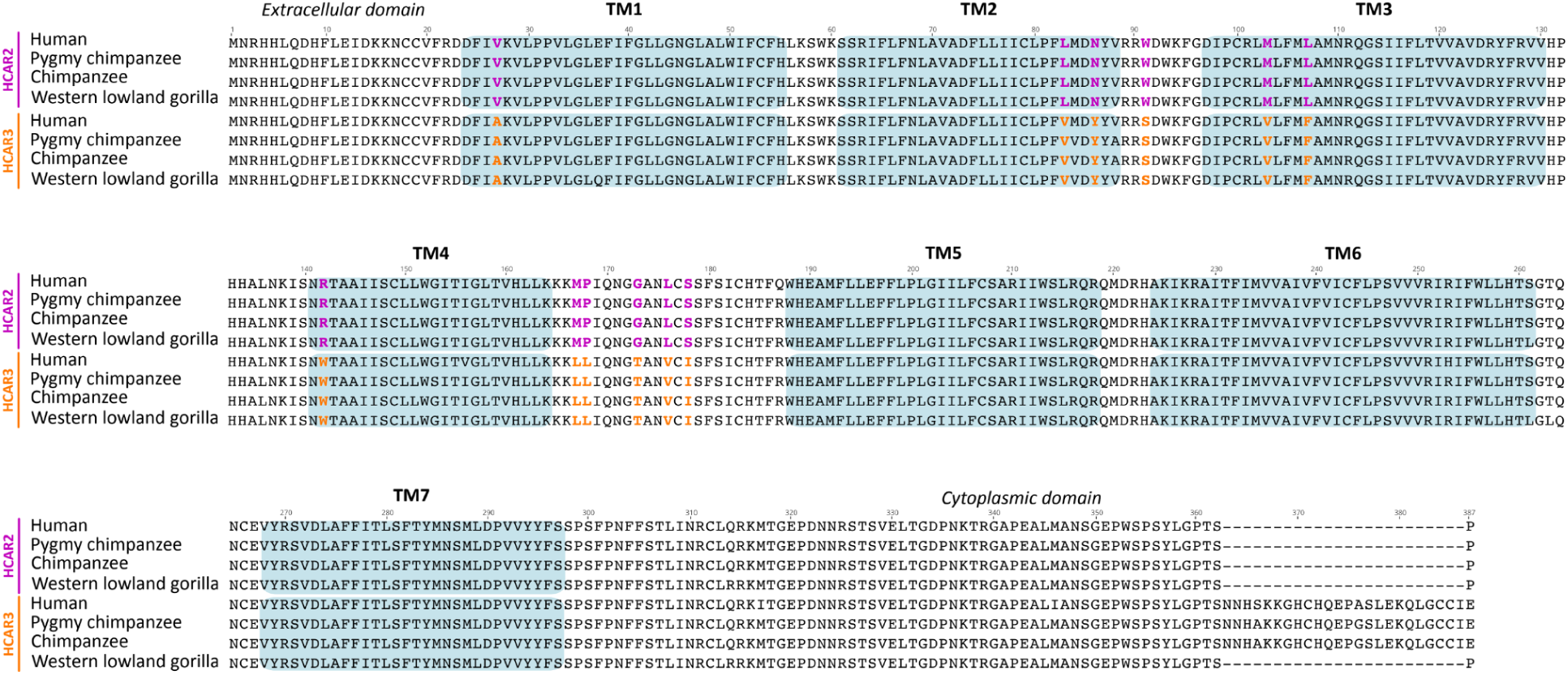
Amino acid alignment of HCAR2 and HCAR3 sequences from human (*Homo sapiens*), chimpanzee (*Pan troglodytes*), pygmy chimpanzee (*Pan paniscus*), and western lowland gorilla (*Gorilla gorilla*). Green shading denotes transmembrane domains. Colored residues highlight the fingerprint positions that unequivocally distinguish HCAR2 from HCAR3.

**Figure 3.**
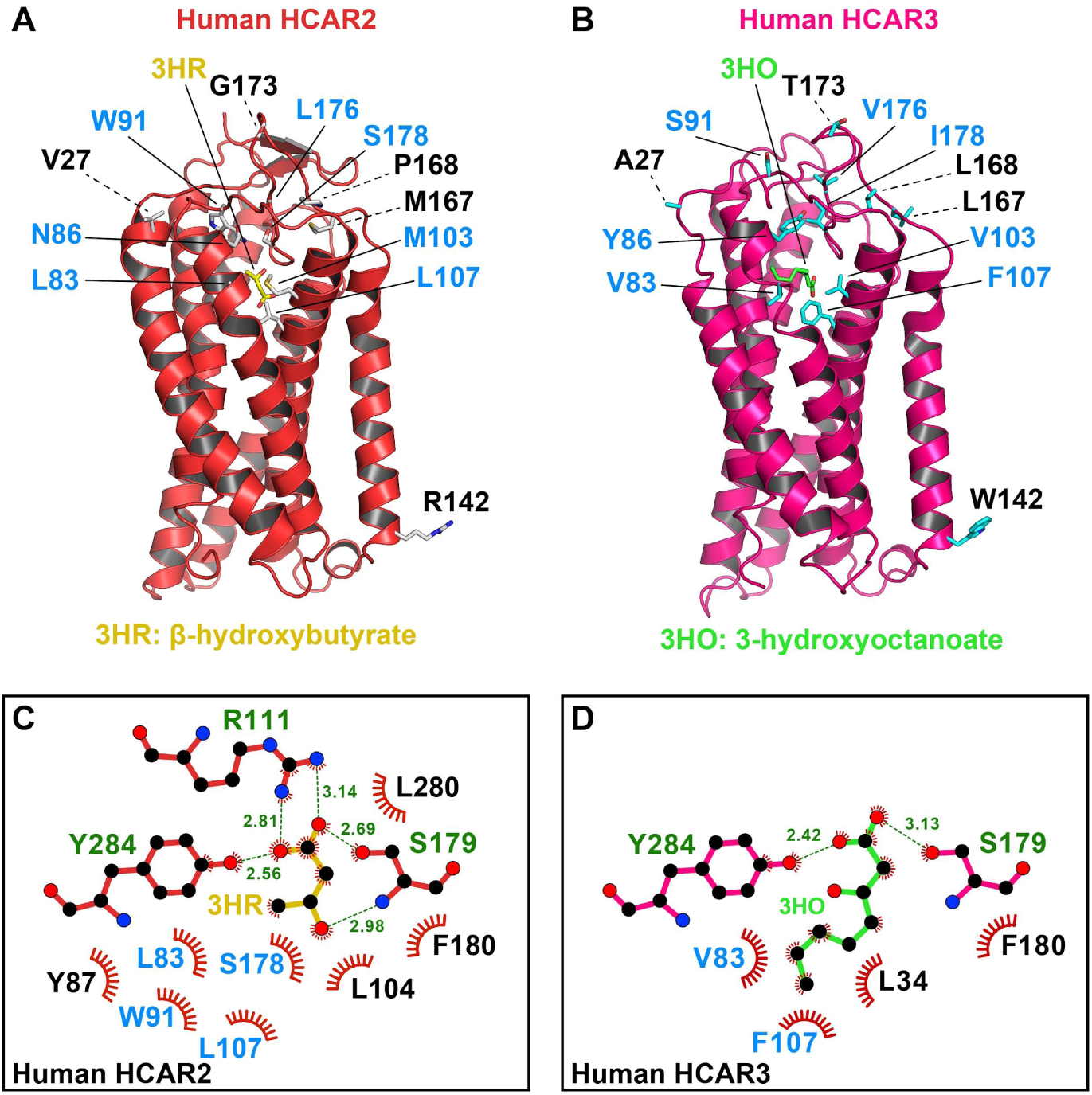
Comparison of distinguishing residues in HCAR2 versus HCAR3. (A) Cartoon representation of human HCAR2 bound to (D)-β-hydroxybutyrate (3HR), shown in stick model with carbon atoms colored yellow (PDB 8J6Q). (B) Cartoon representation of human HCAR3 bound to 3-hydroxyoctanoate (3HO), shown in stick model with carbon atoms colored green (PDB 8JEF). Distinguishing residues between HCAR2 and HCAR3 are shown in stick representation, with those lining the ligand-binding pocket highlighted in blue. (C) Two-dimensional interaction map of (D)-β-hydroxybutyrate (3HR) within the HCAR2 ligand-binding pocket and (D) Two-dimensional interaction map of 3-hydroxyoctanoate (3HO) within the HCAR3 ligand-binding pocket. Residues involved in ligand binding are shown in stick representation, with hydrogen bonds depicted as dashed lines and hydrophobic contacts represented by spoked arcs. Distinguishing residues between HCAR2 and HCAR3 are highlighted in blue.

In addition to these substitutions, a distinctive feature of HCAR3 is the presence of a 24 amino acid insertion in the C-terminal region of the protein in humans, chimpanzees, and pygmy chimpanzees (Fig. 2). Because this insertion is absent in the HCAR3 protein of gorillas, it can be inferred that the insertion arose in the common ancestor of humans and chimpanzees, approximately 6.4 million years ago (Kumar et al. 2017).

### Orangutan-specific Duplications are Recovered in Clades that Exhibit the Functional Signatures of HCAR2 and HCAR3

We utilized the previously defined molecular signatures of the HCAR2 and HCAR3 proteins (Figs. 2 and 3) to assess whether equivalent distinguishing features could be identified in the orangutan-specific duplicates (Fig. 4). Our alignment revealed that ten of the twelve amino acid positions that differentiate HCAR2 from HCAR3 in humans, chimpanzees, pygmy chimpanzees and gorillas also distinguish the orangutan paralogs (Fig. 4). The remaining two positions, residues 86 and 142, associate the orangutan copies more similar to HCAR3 (orange shading) with HCAR2 (gray shading) (Fig. 4). However, the latter residue is unlikely to contribute to ligand-binding differences, as it is located far from the ligand-binding pocket (Fig. 3). Interestingly, the orangutan duplicates most similar to HCAR3 lack the 24 amino acid insertion found in the HCAR3 sequences of humans, chimpanzees and pygmy chimpanzees (Figs. 2 and 4). Thus, based on these diagnostic amino acid residues, we hypothesize that the orangutan-specific duplicates may functionally resemble their human counterparts, despite not being 1:1 orthologs. Supporting the hypothesis of independent evolutionary origins, cAMP inhibition assays assessing HCAR3 responsiveness to D-phenyllactic acid, a potent HCAR3 agonist, revealed two separable functional groups (Peters et al. 2019). One group comprised the 1:1 HCAR3 orthologs from human, chimpanzee, and gorilla, whereas the second consisted of the independently originated orangutan HCAR3 (Peters et al. 2019).

**Figure 4.**
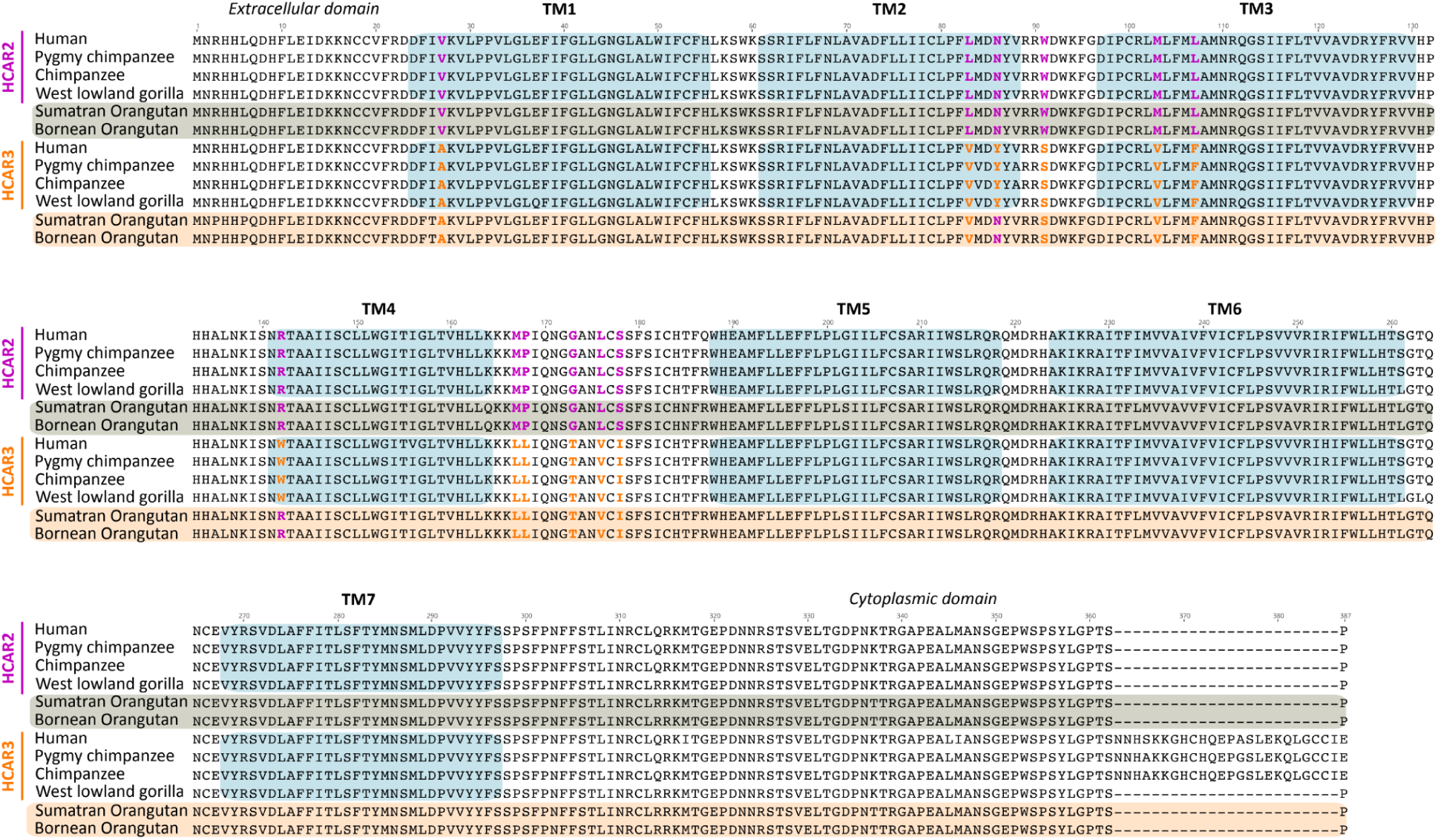
Amino acid alignment of HCAR2 and HCAR3 sequences from human (*Homo sapiens*), chimpanzee (*Pan troglodytes*), pygmy chimpanzee (*Pan paniscus*), and western lowland gorilla (*Gorilla gorilla*), along with the duplicate copies from Sumatran and Bornean orangutans (*Pongo abelii* and *Pongo pygmaeus*, respectively). TM indicates transmembrane domains. Colored residues highlight the fingerprint positions that associate each orangutan duplicate with either HCAR2 or HCAR3.

### Lesser Apes have a More Dynamic Evolutionary Trend in Comparison to Great Apes

In lesser apes, the gene turnover rate is more dynamic than in great apes, leading to a more complex evolutionary history (Fig. 1). Our phylogenetic analysis recovered a well-supported clade comprising three species (Fig. 1, green branches): the agile gibbon (*Hylobates agilis*) and the northern white-cheeked gibbon (*Nomascus leucogenys*), each with a single gene copy, and the siamang (*Symphalangus syndactylus*), which possesses two copies, one of which falls within this clade (Fig. 1). By reconciling this gene tree with the species phylogeny of lesser apes (Goodman et al. 1979; Shao et al. 2023), we infer that this gene lineage was present in the common ancestor of the group and was subsequently lost in the species of the genus *Hoolock*. Based on twelve previously identified diagnostic fingerprint residues, the sequences within the green clade are predicted to be functionally equivalent to HCAR2 (Fig. 5). Specifically, eleven of the twelve positions match the HCAR2 profile, with only a single mismatch at position 27. This residue corresponds to the HCAR3 signature and is located far from the ligand-binding pocket, making it unlikely to influence ligand interactions (Fig. 5). Interestingly, the two gibbon species that retained a single HCAR gene preserved the copy that is functionally equivalent to HCAR2 (Fig. 5).

**Figure 5.**
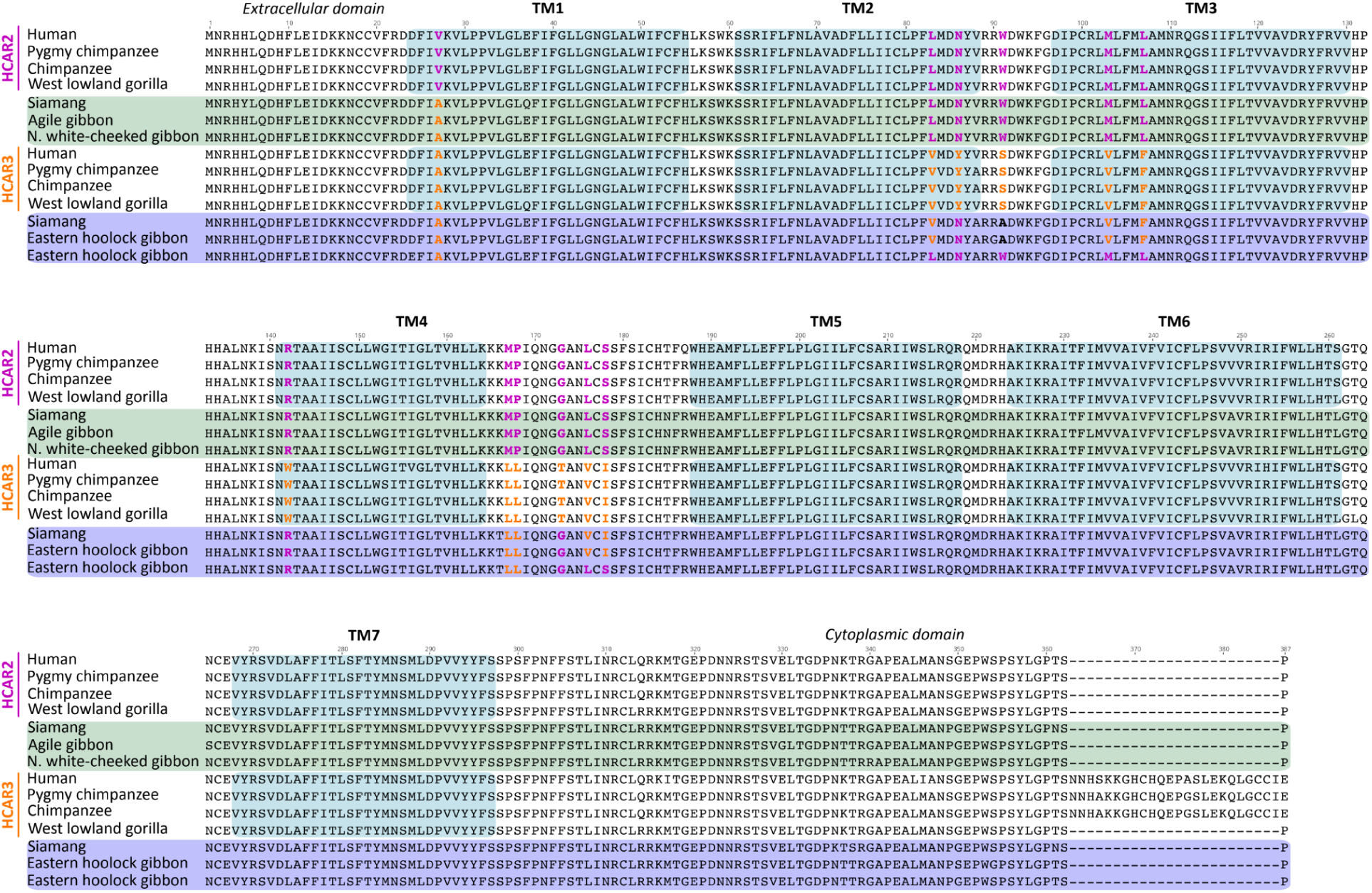
Amino acid alignment of HCAR2 and HCAR3 sequences from human (*Homo sapiens*), chimpanzee (*Pan troglodytes*), pygmy chimpanzee (*Pan paniscus*), and western lowland gorilla (*Gorilla gorilla*), along with the duplicate copies from siamang (*Symphalangus syndactylus*) and hoolock gibbon (*Hoolock leuconedys*). Included are also the single copies from the agile gibbon (*Hylobates agilis*) and the northern white-cheeked gibbon (*Nomascus leucogenys*). TM indicates transmembrane domains. Colored residues highlight the fingerprint positions that associate each lesser ape protein with either HCAR2 or HCAR3.

In the other lesser ape clade (Fig. 1, blue branches), we recovered the second copy of the siamang gene and two copies from the hoolock gibbon (*Hoolock leuconedys*). This topology suggests a secondary duplication event that is retained only in the genus *Hoolock*. Functional predictions based on the diagnostic residues indicate that the siamang copy has eight positions that match HCAR3, three match those of HCAR2, and one does not match any paralog (Fig. 5). This pattern makes functional sense as the siamang copy recovered in the other clade has HCAR2 functional identity (Fig. 5). Thus, according to our analyses, the siamang would have the functional equivalent of HCAR2 and HCAR3 in its genome. In the case of the hoolock gibbon, the two gene copies exhibit contrasting profiles. One copy shares eight HCAR3 diagnostic positions, three HCAR2 positions, and one ambiguous site, closely mirroring the pattern observed in the siamang HCAR3-like copy. The other copy displays nine matches with HCAR2 and three with HCAR3 (Fig. 5), suggesting functional divergence between the paralogs. Thus, similar to the case of the siamang, the hoolock gibbon has copies functionally equivalent to HCAR2 and HCAR3 in its genome.

For this primate lineage, it has been proposed that the siamang HCAR2 and HCAR3 receptors have experienced a relaxation of evolutionary constraint (Peters et al. 2019). This interpretation was based on d_N_/d_S_ analyses showing that, although the estimated ω falls within the range typically associated with purifying selection, the likelihood of this model was not significantly different from that of a model in which ω is fixed at 1. However, it is important to note that reliable inference of selective regimes depends strongly on data requirements, particularly the number of species included, which directly affects statistical power (Anisimova et al. 2001; McBee et al. 2015). The authors further argued that, in contrast to other HCAR orthologs, siamang HCAR2 and HCAR3 exhibit less clearly differentiated functional profiles, which they interpreted as evidence of reduced functional conservation. In our analyses, however, the reduced functional differentiation reported by the authors may instead reflect the fact that the two siamang copies are not clearly differentiated at the sequence level. Specifically, the copy recovered within the green clade retains 11 of the 12 diagnostic positions consistent with the HCAR2 profile, with only a single site (position 27), located far from the binding pocket, matching the HCAR3 profile. In contrast, the copy recovered within the blue clade shows a more ambiguous diagnostic signature as eight positions match the HCAR3 profile, three positions match the HCAR2 profile, and one site (position 91) does not correspond to either profile. Importantly, although the authors interpreted these patterns by comparison with other ape lineages, such comparisons cannot be made directly because the siamang sequences are not 1:1 orthologs of HCAR2 and HCAR3 in other ape species, as originally claimed. Rather, the siamang copies appear to have followed distinct evolutionary trajectories, most likely because they arose from independent duplication events. Thus, the apparent lack of functional differentiation in siamang relative to other apes may be better explained by differences in evolutionary origin and lineage-specific history, rather than by a relaxation of evolutionary constraint.

### Primate Species that Retained the Ancestral Condition of a Single-Copy Gene Possess a Gene Functionally Equivalent to HCAR2

As we already know, only apes possess duplicated copies corresponding to the HCAR2 and HCAR3 genes, which encode receptors with distinct ligand specificities. HCAR2 binds nicotinic acid and (D)-β-hydroxybutyrate with high affinity, mediating antilipolytic and antiinflammatory responses upon ligand activation. In contrast, HCAR3, which binds 3-hydroxyoctanoic acid, plays a key role in regulating lipid metabolism by modulating lipolysis in adipose tissue. In the case of all other primate groups, i.e., Old World Monkeys, New World monkeys, tarsiers, and strepsirhines, they have only a single copy of a gene that encodes an HCAR2/3 receptor (Fig. 1). Under this scenario, the key question is to which of the paralogs (HCAR2 or HCAR3) is the single-copy gene (HCAR2/3) in non-ape primates functionally equivalent? To address this question, we employed the twelve diagnostic residues previously identified by comparing the four species that possess, from an evolutionary perspective, HCAR2 and HCAR3 genes (Figs. 2 and 3). As shown in Figure 6, all non-ape primate species have conserved the full set of seven amino acid residues that are located in close proximity to the ligand-binding site (Fig. 3), suggesting that these receptors are likely functionally equivalent to HCAR2. In contrast, limited variation is observed among residues positioned distal to the receptor binding site (Fig. 6).

**Figure 6.**
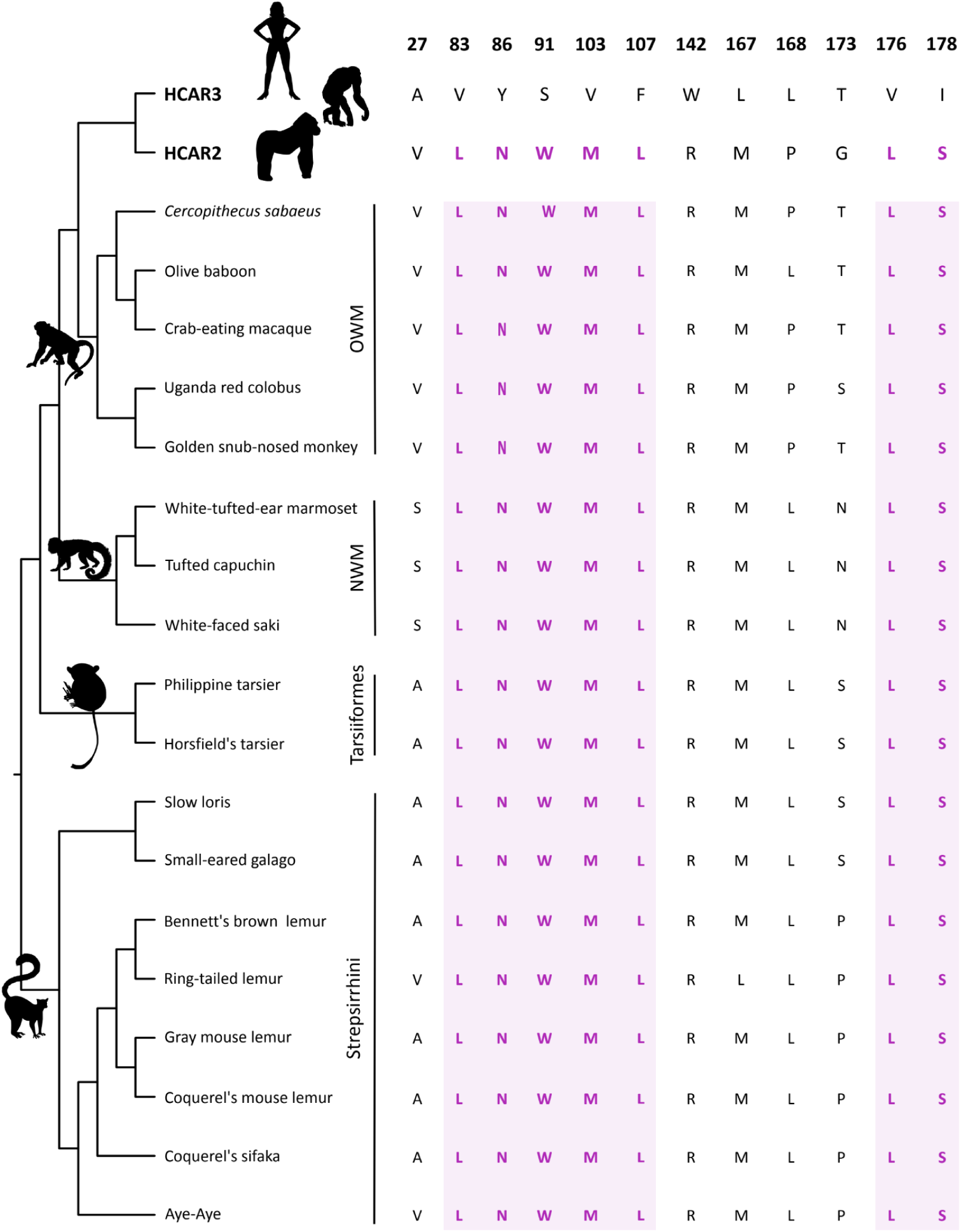
Conservation pattern of the twelve diagnostic amino acid residues previously identified on HCAR2 and HCAR3 in non-ape primate species. Residues highlighted in pink represent amino acids located in close proximity to the ligand-binding site. In contrast, non-highlighted residues are positioned distal to the binding site and are therefore unlikely to be directly involved in ligand interaction. For simplicity, duplicated copies from orangutans and gibbons were omitted. Silhouette images were obtained from PhyloPic (http://phylopic.org/).

In Old World monkeys, the sister group of apes, eleven of the diagnostic residues are identical to those of HCAR2 (Fig. 6). The only exception is position 173, which corresponds to the HCAR3 variant in all species except the Uganda red colobus, where a serine is present (Fig. 6). This variation is unlikely to have functional consequences, as our structural analyses indicate that this residue is located far from the ligand-binding pocket (Fig. 6 and Supplementary Figure S1). It is also interesting to note that species belonging to the subfamily Colobinae have a deletion of eleven residues in the C-terminal region of the protein (Supplementary Figure S1). Thus, based on the twelve diagnostic residues previously identified, the protein encoded by the HCAR2/3 gene in Old World monkeys is likely functionally equivalent to HCAR2.

**Supplementary figure S1.**
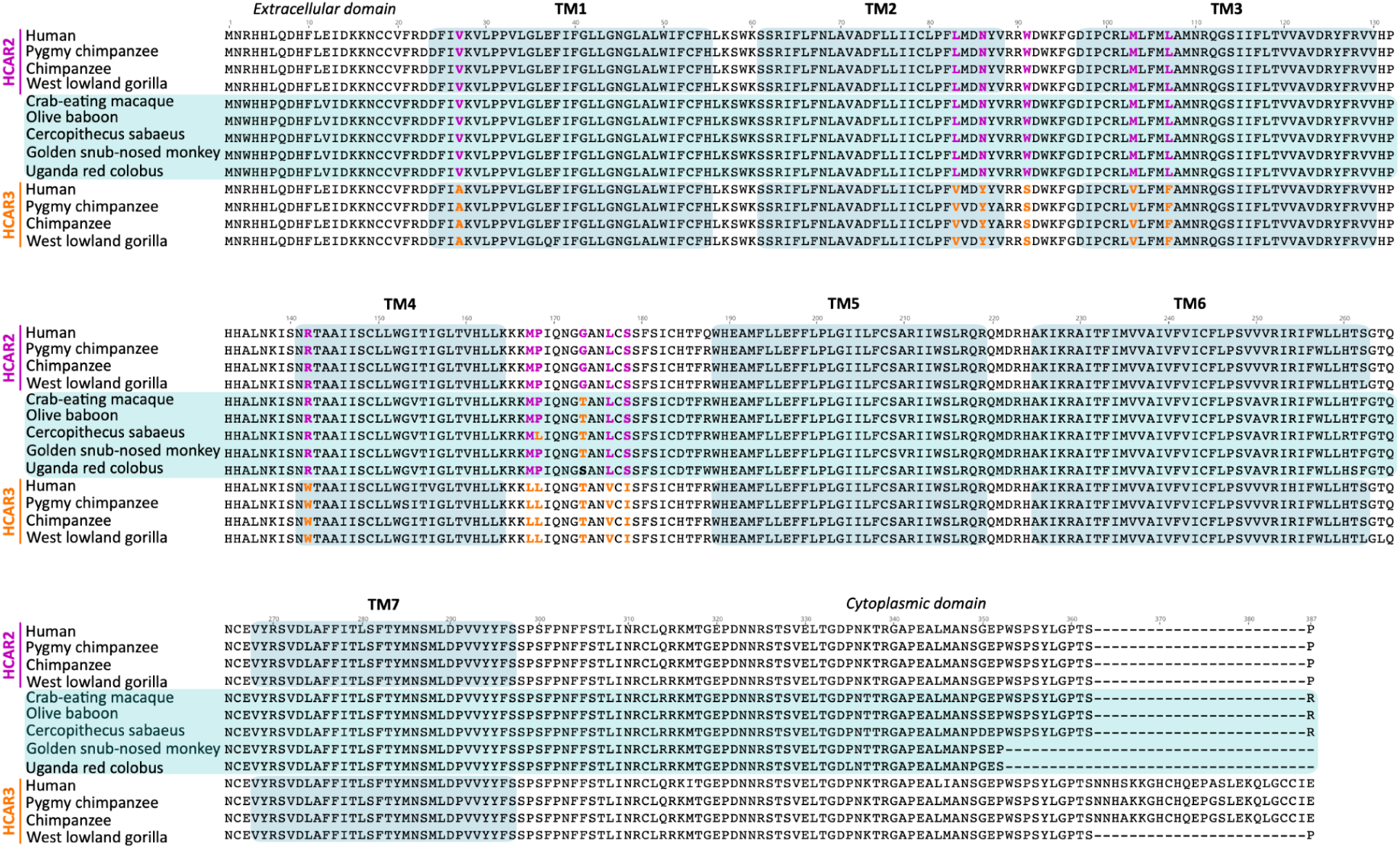
Amino acid alignment of HCAR2 and HCAR3 sequences from human (*Homo sapiens*), chimpanzee (*Pan troglodytes*), pygmy chimpanzee (*Pan paniscus*), and western lowland gorilla (*Gorilla gorilla*), along with the single copy gene (HCAR2/3) from the crab-eating macaque (*Macaca fascicularis*), Olive baboon (*Papio anubis*), *Cercopithecus sabaeus*, Uganda red colobus (*Piliocolobus tephrosceles*), and golden snub-nosed monkey (*Rhinopithecus roxellana*). TM indicates transmembrane domains. Colored residues highlight the diagnostic alignment positions.

In New World monkeys, the pattern of diagnostic positions is broadly similar to that observed in Old World monkeys. Specifically, nine of the diagnostic residues match the HCAR2 protein profile, while one corresponds to the HCAR3 profile (position 168, Fig. 6 and Supplementary Figure S2). Interestingly, two diagnostic positions in this group do not match either HCAR2 or HCAR3 (Fig. 6 and Supplementary Figure S2). At position 27, where HCAR2 has a valine and HCAR3 an alanine, New World monkeys instead exhibit a serine (Fig. 6 and Supplementary Figure S2). Similarly, at position 173, HCAR2 contains a glycine and HCAR3 a threonine, whereas New World monkeys harbor an asparagine at this site (Fig. 6 and Supplementary Figure S2). These discordances are unlikely to have functional consequences, as our structural analyses indicate that the affected residues are located far from the ligand-binding pocket (Fig. 3). Taken together, these observations suggest that the protein encoded by the gene corresponding to the pre-duplication state (HCAR2/3) in New World monkeys is most likely functionally equivalent to HCAR2.

**Supplementary figure S2.**
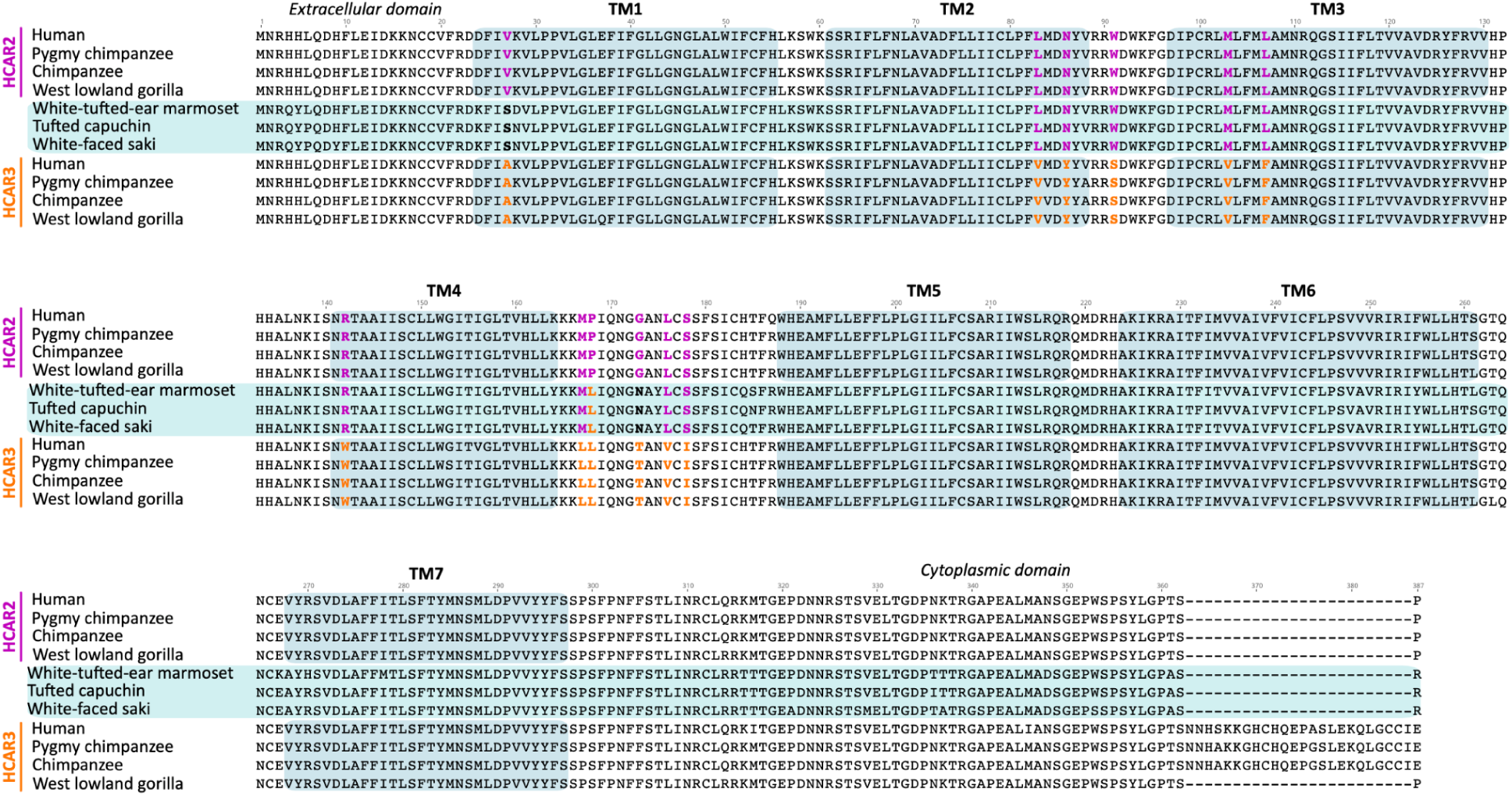
Amino acid alignment of HCAR2 and HCAR3 sequences from human (*Homo sapiens*), chimpanzee (*Pan troglodytes*), pygmy chimpanzee (*Pan paniscus*), and western lowland gorilla (*Gorilla gorilla*), along with the single copy gene (HCAR2/3) from the white-tufted-ear marmoset (*Callithrix jacchus*), tufted capuchin (*Sapajus apella*), and white-faced saki (*Pithecia pithecia*). TM indicates transmembrane domains. Colored residues highlight the diagnostic alignment positions.

In tarsiers, small primates from Southeast Asia known for their large eyes and powerful leaping legs, the pattern supports what has been observed in other primate groups that retain the ancestral condition of a single-copy gene (HCAR2/3), i.e., that it would be functionally equivalent to HCAR2 (Fig. 6 and Supplementary Figure S3). In this primate group, nine diagnostic positions match the HCAR2 protein profile, while two correspond to the HCAR3 profile (Fig. 6 and Supplementary Figure S3). Interestingly, similar to the situation observed in New World monkeys, the diagnostic position 173 does not match either HCAR2 or HCAR3 protein profiles (Fig. 6 and Supplementary Figure S3). While HCAR2 has a glycine and HCAR3 a threonine, tarsiers possess a serine at this site (Fig. 6 and Supplementary Figure S3). Thus, the pattern observed in tarsiers provides further support for the conclusion that, in primate species possessing a single copy of the gene, this copy is functionally equivalent to HCAR2.

Finally, in strepsirrhines, a primate group characterized by features such as a moist rhinarium, a toothcomb, and a predominantly nocturnal lifestyle, the observed pattern supports the same conclusion, namely that the single-copy gene present in this group would be functionally equivalent to HCAR2 (Fig. 6 and Supplementary Figure S4). In this primate group, nine diagnostic positions match the HCAR2 protein profile, whereas two positions (positions 27 and 168) correspond to the HCAR3 profile (Fig. 6 and Supplementary Figure S4). As observed in tarsiers and New World monkeys, position 173 does not match either the HCAR2 or HCAR3 profiles. HCAR2 has a glycine and HCAR3 a threonine at this site, while strepsirrhines possess either a proline or a serine (Fig. 6 and Supplementary Figure S4). At position 167, most species match the HCAR2 profile, however, the ring-tailed lemur (*Lemur catta*) possesses a leucine matching the HCAR3 profile (Fig. 6 and Supplementary Figure S4). Finally, at position 27, two species (aye-aye, *Daubentonia madagascariensis*, and ring-tailed lemur, *Lemur catta*) match the HCAR2 profile, whereas the remaining two species correspond to the HCAR3 profile (Fig. 6 and Supplementary Figure S4).

**Supplementary figure S3.**
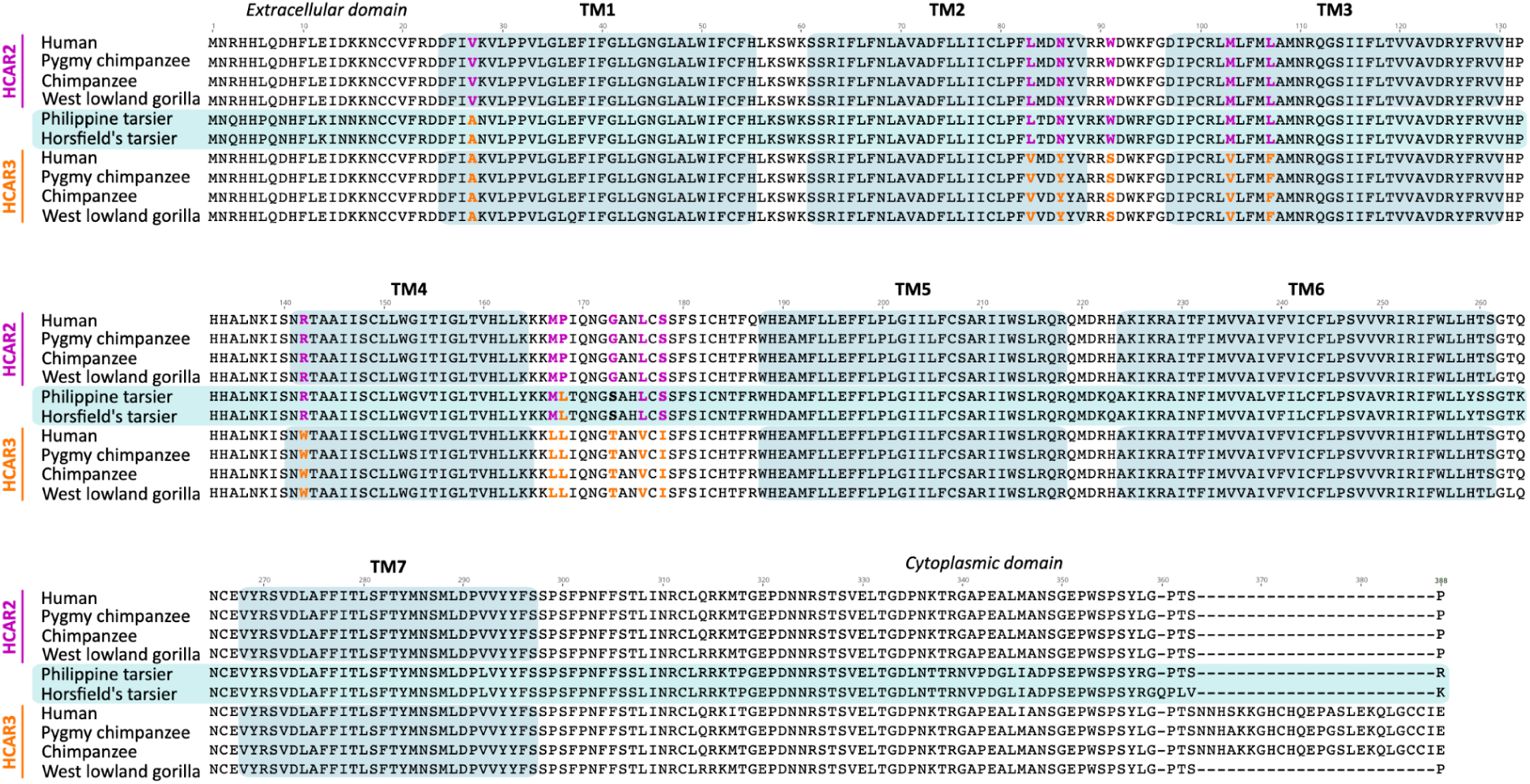
Amino acid alignment of HCAR2 and HCAR3 sequences from human (*Homo sapiens*), chimpanzee (*Pan troglodytes*), pygmy chimpanzee (*Pan paniscus*), and western lowland gorilla (*Gorilla gorilla*), along with the single copy gene (HCAR2/3) from the Philippine tarsier (*Carlito syrichta*), and Horsfield’s tarsier (*Cephalopachus bancanus*). TM indicates transmembrane domains. Colored residues highlight the diagnostic alignment positions.

**Supplementary figure S4.**
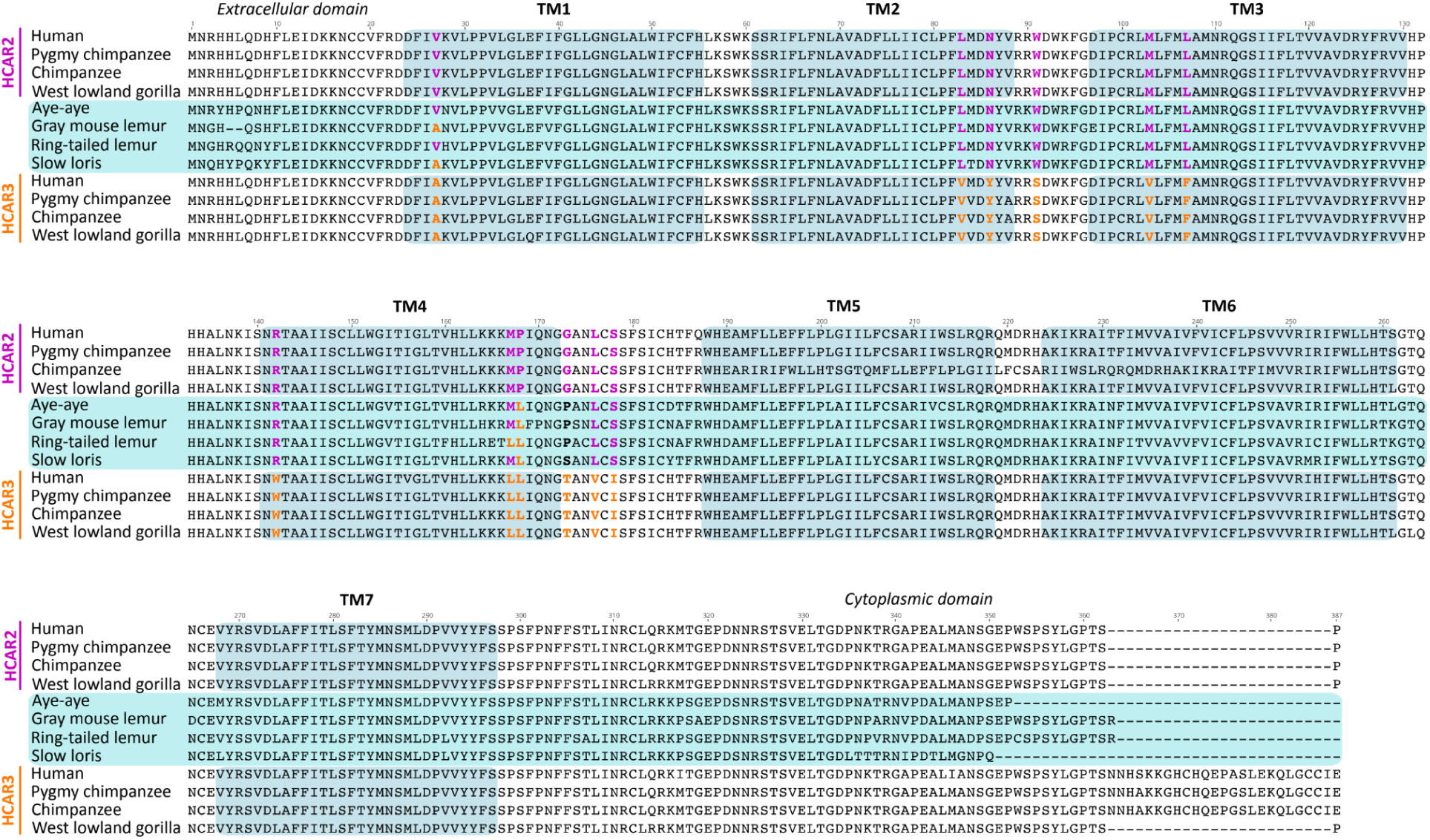
Amino acid alignment of HCAR2 and HCAR3 sequences from human (*Homo sapiens*), chimpanzee (*Pan troglodytes*), pygmy chimpanzee (*Pan paniscus*), and western lowland gorilla (*Gorilla gorilla*), along with the single copy gene (HCAR2/3) from the aye-aye (*Daubentonia madagascariensis*), gray mouse lemur (*Microcebus murinus*), ring-tailed lemur (*Lemur catta*), and slow loris (*Nycticebus coucang*). TM indicates transmembrane domains. Colored residues highlight the diagnostic alignment positions.

### Structural Analysis of the HCAR2/3 Receptors Further Supports Functional Equivalence with HCAR2

Previously, by comparing residues that unambiguously distinguish HCAR2 from HCAR3, we concluded that non-ape species possessing a single-copy gene encode a receptor that would be functionally equivalent to HCAR2 (Fig. 6). To strengthen this conclusion further, we performed three-dimensional structural analyses by generating structural models of non-ape primate HCAR2/3 receptors and subsequently modeling their complexes with either (D)-β-hydroxybutyrate or 3-hydroxyoctanoate (Supplementary Figure S5, S6, and Fig. 7A and 7B). As expected, the predicted structures, with no ligands, of all HCAR2/3 receptors were highly similar to both human HCAR2 and human HCAR3 (Figs. 7A). However, the RMSD values calculated from superimposed Cα coordinates showed that the HCAR2/3 models were significantly more similar to human HCAR2 (0.801 ± 0.173 Å) than to human HCAR3 (1.336 ± 0.082 Å) (Fig. 7A) (Chothia and Lesk 1986; Sali and Blundell 1993). Moreover, models of non-ape primate HCAR2/3 receptors exhibited a better fit when bound to (D)-β-hydroxybutyrate than to 3-hydroxyoctanoate, i.e., in each HCAR2/3 model, the spatial orientation of (D)-β-hydroxybutyrate closely matched that observed in human HCAR2 (Fig. 8), including the conserved proximity of several amino acids within the ligand-binding pocket (Fig. 9). In contrast, in all HCAR2/3 models, the spatial disposition of 3-hydroxyoctanoate showed an orientation opposite to that found in human HCAR3 and adopted a different axial alignment (Supplementary Figure S7). These spatial dispositions are consistent with AlphaFold3 models of 3-hydroxyoctanoate bound to human HCAR2 (Supplementary Figure S8A and B) and of (D)-β-hydroxybutyrate bound to HCAR3 (Supplementary Figure S8C and D), strongly suggesting that non-ape primate HCAR2/3 receptors are functionally equivalent to ape HCAR2. Of note, the latter models also revealed that none of the distinguishing residues directly stabilized the bound ligands, suggesting that ligand binding can be maintained through alternative networks of interactions and that ligand specificity relies on additional features beyond the identity of ligand-binding pocket residues.

**Figure 7.**
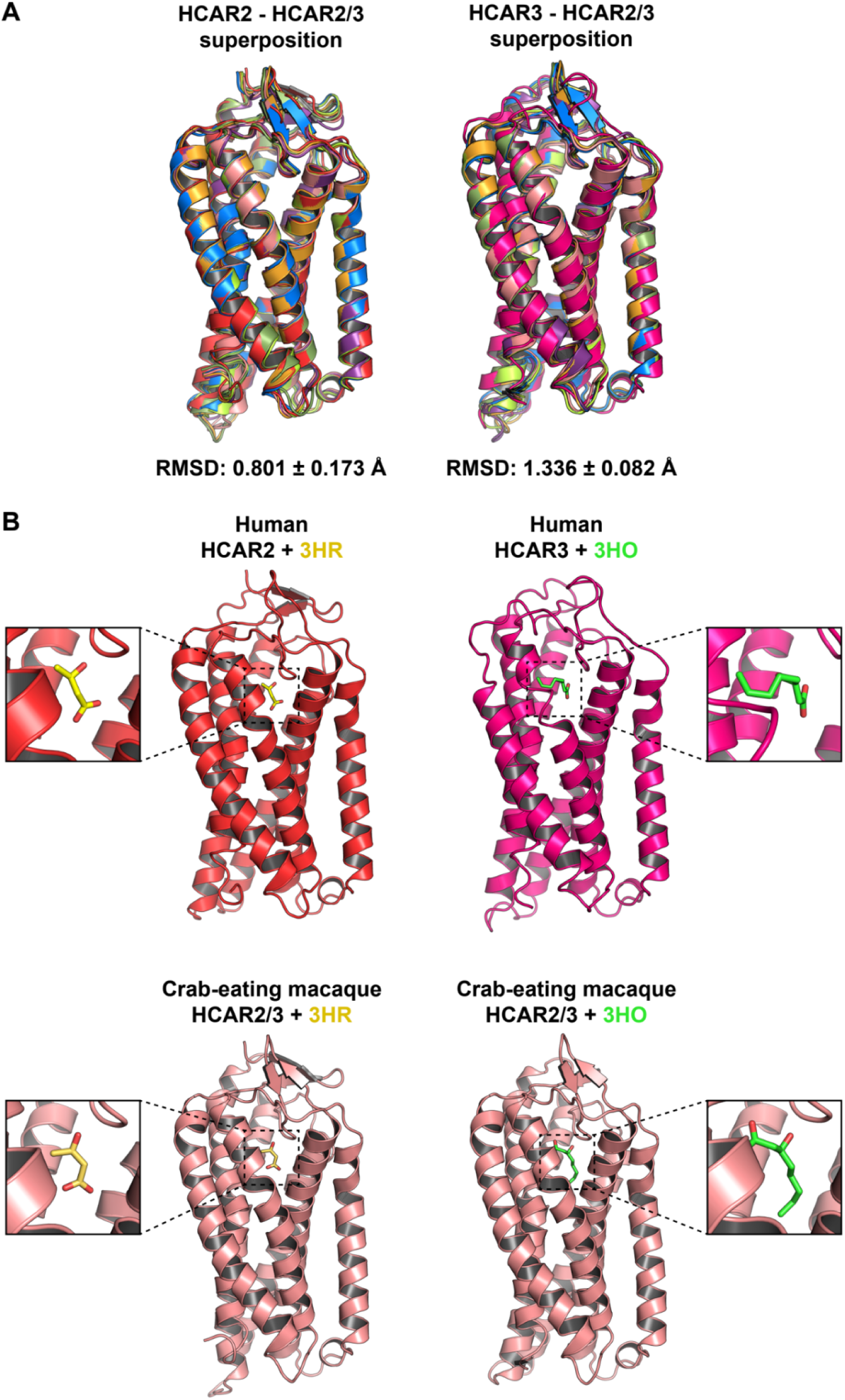
Comparison of HCAR2/3 structural models with human HCAR2 and human HCAR3. (A) Cartoon representation of the superposition of the structural models of human HCAR2 (tv red) or human HCAR3 (hot pink) with the HCAR2/3 structural models from non-ape primates (White-tufted-ear marmoset (*Callithrix jacchus*): bright orange; Crab-eating macaque (*Macaca fascicularis*): salmon; Golden snub-nosed monkey (*Rhinopithecus roxellana*): violet purple; Philippine tarsier (*Carlito syrichta*): smudge green; Coquerel’s sifaka (*Propithecus coquereli*): limon yellow; Ring-tailed lemur: marine blue). (B) Cartoon representation of the indicated HCAR2 (tv red), HCAR3 (hot pink), and HCAR2/3 (salmon) structural models bound to (D)-β-hydroxybutyrate (3HR) or to 3-hydroxyoctanoate (3HO). Zoomed views highlight the spatial orientation of the ligands within the binding pocket. Note the closely parallel orientation of 3HR in HCAR2 and HCAR2/3 models, and the closely perpendicular orientation of 3HO in HCAR3 and HCAR2/3 models.

**Figure 8.**
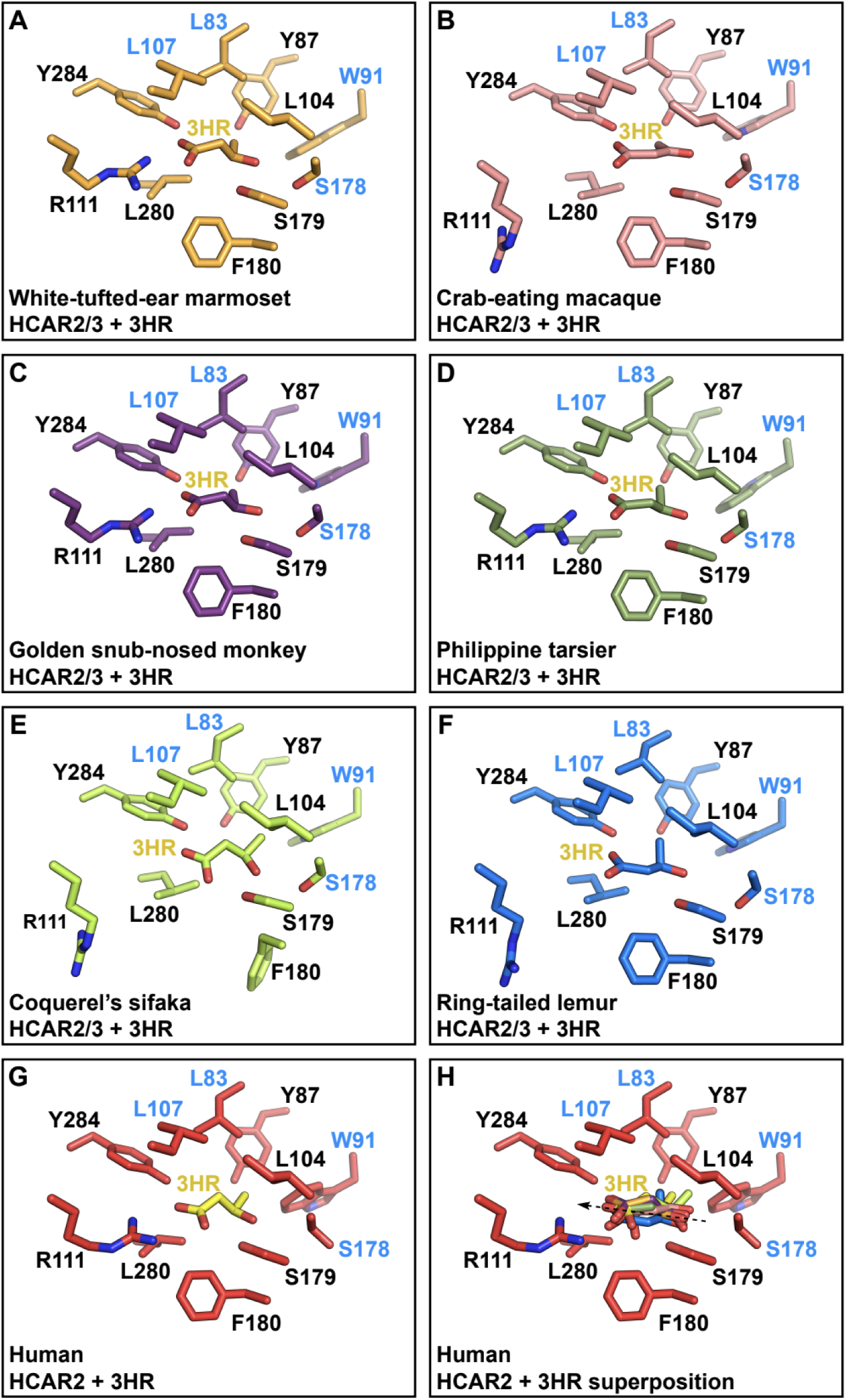
Comparison of the ligand-binding pocket of HCAR2/3 models bound to (D)-β-hydroxybutyrate. (A-F) Stick representation of HCAR2/3 ligand binding pocket amino acid residues of the indicated non-ape primate species bound to β-hydroxybutyrate (3HR). (G) For comparison, human HCAR2 ligand-binding pocket amino acid residues are shown bound to 3HR (PDB 8J6Q). (H) Comparison of the orientation of each 3HR shown in stick representation after the superposition of all HCAR2/3 models (A-F) with human HCAR2 (G). The dashed arrow depicts the overall orientation of superposed 3HR ligands. Distinguishing residues between HCAR2 and HCAR3 are highlighted in blue.

**Figure 9.**
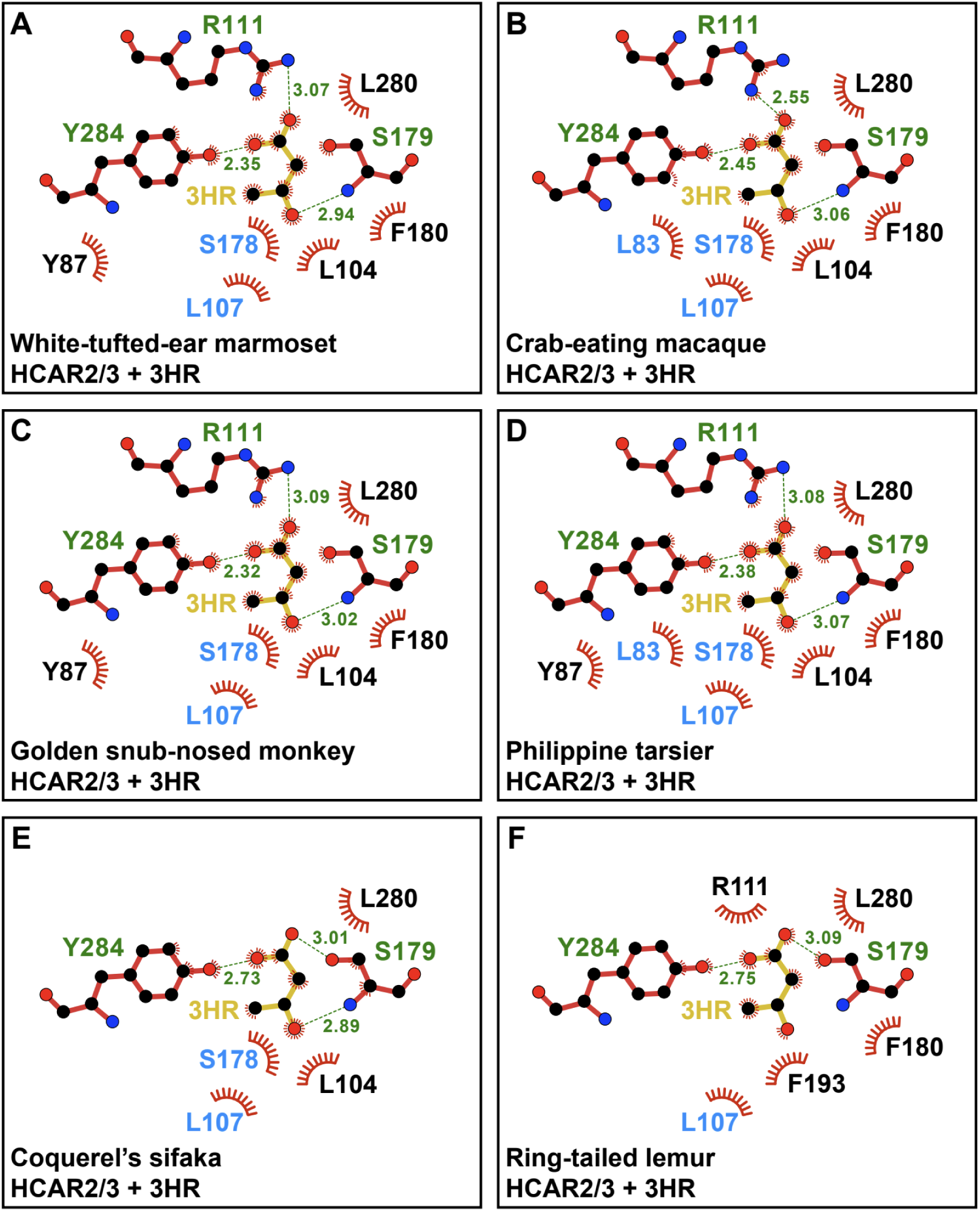
Comparison of the ligand-binding pocket of HCAR2/3 bound to β-hydroxybutyrate using LigPlot^+^. (A-F) Two-dimensional, schematic representation of the interactions of (D)-β-hydroxybutyrate (3HR) with the indicated amino acid residues within the ligand-binding pocket of the HCAR2/3 models of the indicated species. Distinguishing residues between HCAR2 and HCAR3 are highlighted in blue.

**Supplementary Figure S5.**
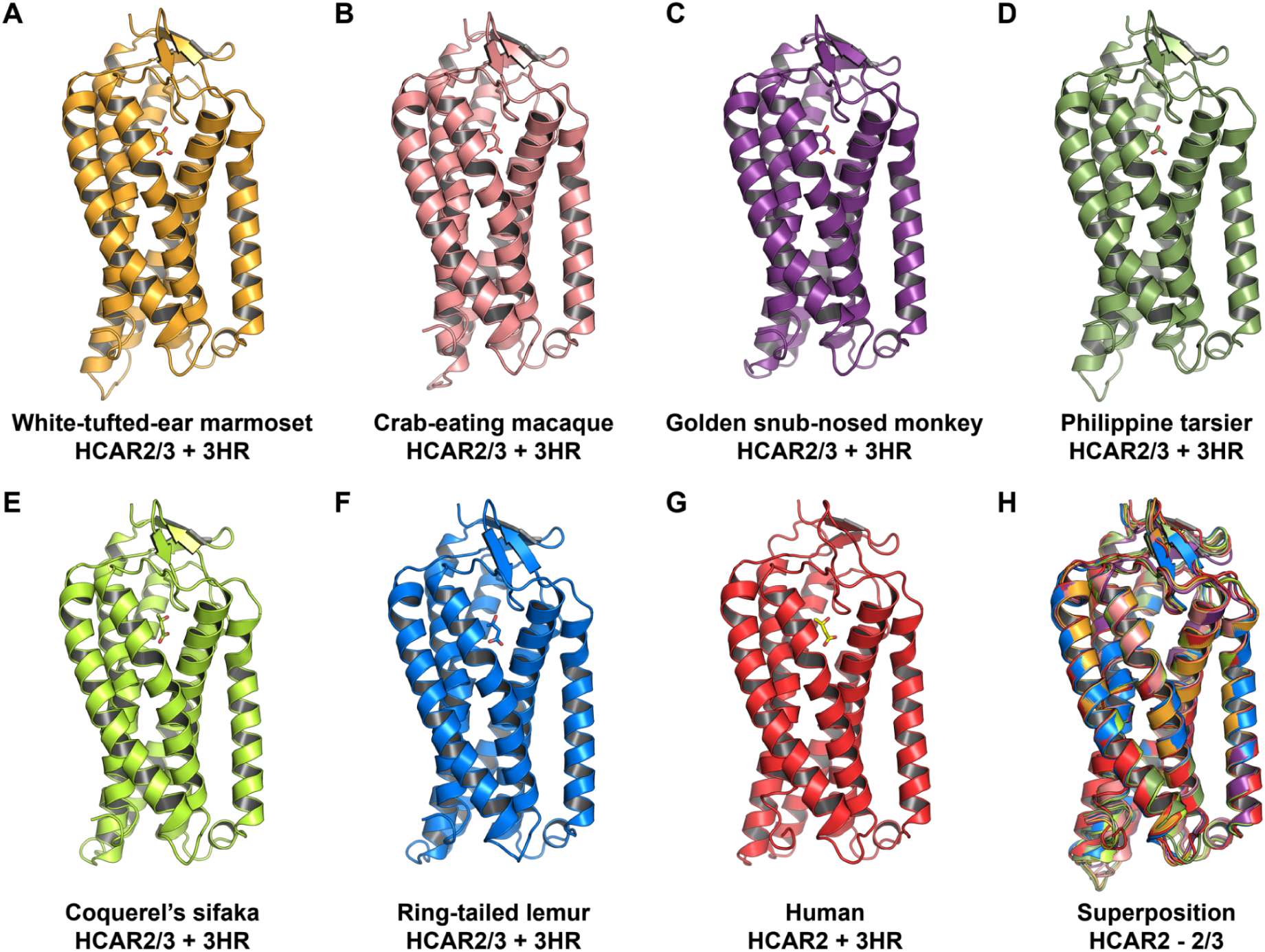
Structural models of non-ape primate HCAR2/3 receptor bound to (D)-β-hydroxybutyrate generated by AlphaFold3. (A-F) Cartoon representations of HCAR2/3 receptors of White-tufted-ear marmoset (*Callithrix jacchus*) (A), Crab-eating macaque (*Macaca fascicularis*) (B), Golden snub-nosed monkey (*Rhinopithecus roxellana*) (C), Philippine tarsier (*Carlito syrichta*) (D), Coquerel’s sifaka (*Propithecus coquereli*) (E), Ring-tailed lemur (*Lemur catta*) (F) bound to (D)-β-hydroxybutyrate (3HR), shown in stick representation. (G) For comparison, the cartoon representation of human HCAR2 is shown bound to 3HR, depicted in stick representation (PDB 8J6Q). (H) Cartoon representation of the Cα coordinate superposition of the models shown in A to G that showed a RMSD of 0.801 ± 0.173 Å.

**Supplementary Figure S6.**
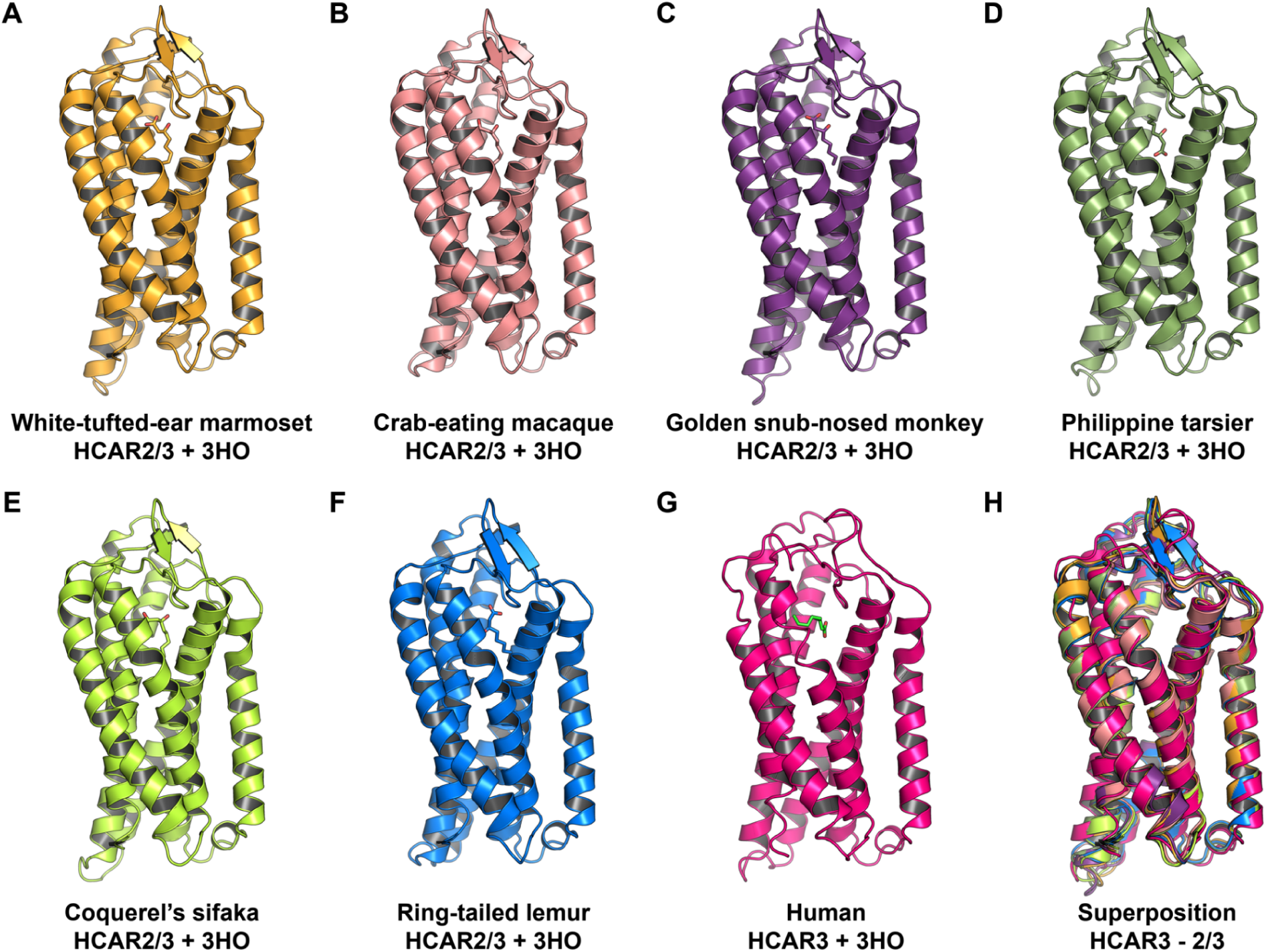
Structural models of non-ape primate HCAR2/3 bound to 3-hydroxyoctanoate generated by AlphaFold3. (A-F) Cartoon representations of HCAR2/3 receptors of White-tufted-ear marmoset (*Callithrix jacchus*) (A), Crab-eating macaque (*Macaca fascicularis*) (B), Golden snub-nosed monkey (*Rhinopithecus roxellana*) (C), Philippine tarsier (*Carlito syrichta*) (D), Coquerel’s sifaka (*Propithecus coquereli*) (E), Ring-tailed lemur (*Lemur catta*) (F) bound to 3-hydroxyoctanoate (3HO), shown in stick representation. (**G**) For comparison, the cartoon representation of human HCAR2 is shown bound to 3HO, depicted in stick representation (PDB 8JEF). (**H**) Cartoon representation of the Cα coordinates superposition of the models shown in A to G that showed a RMSD of 1.336 ± 0.082 Å.

**Supplementary Figure S7.**
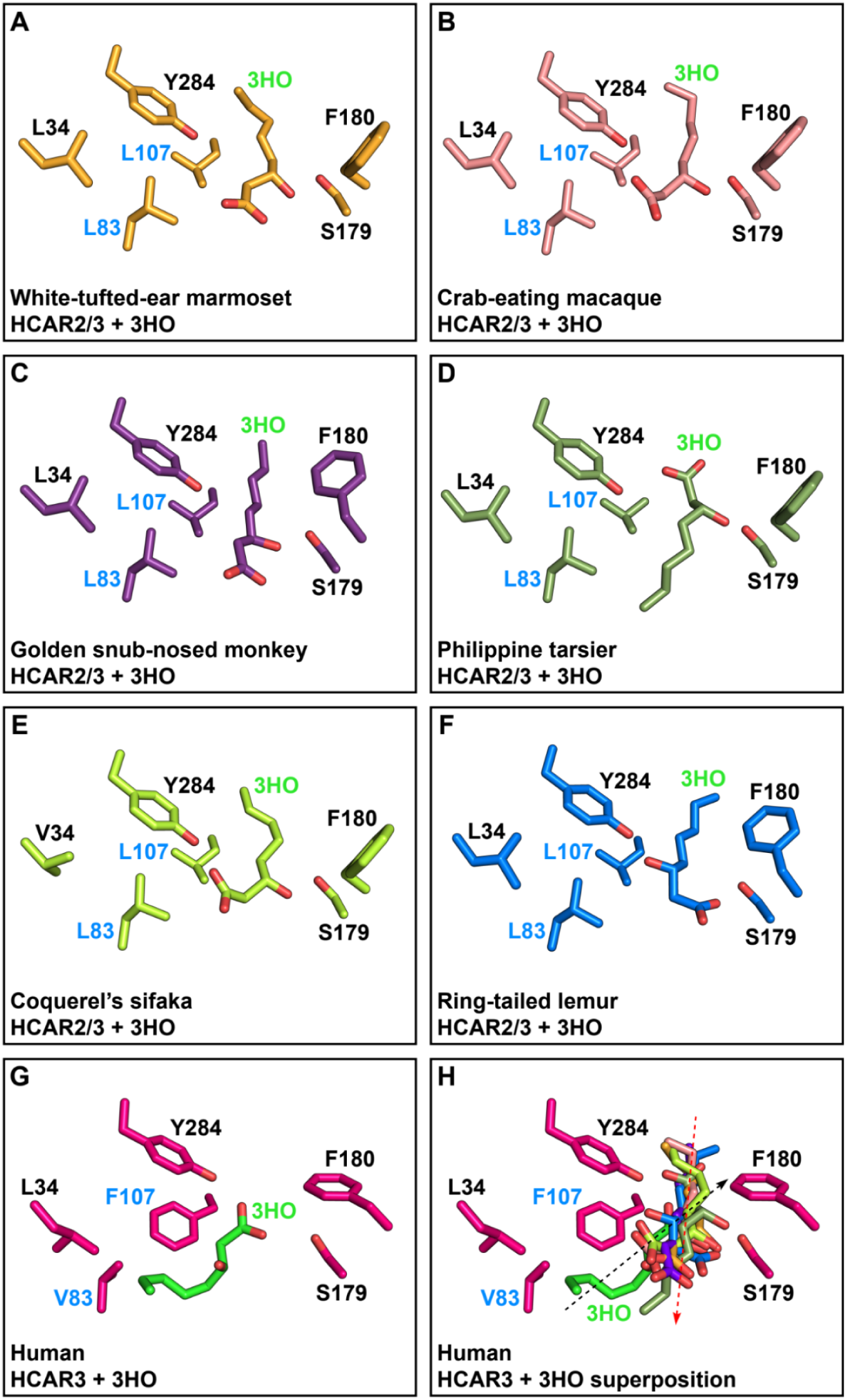
Comparison of the ligand-binding pocket of HCAR2/3 models bound to 3-hydroxyoctanoate. (A-F) Stick representation of HCAR2/3 ligand binding pocket amino acid residues of the indicated non-ape primate species bound to 3-hydroxyoctanoate (3HO). (G) For comparison, human HCAR3 ligand-binding pocket amino acid residues are shown bound to 3HO (PDB 8JEF). (H) Comparison of the orientation of each 3HO shown in stick representation after the superposition of all HCAR2/3 models (A-F) with human HCAR3 (G). The dashed black arrow depicts the orientation of 3HO within the ligand-binding pocket of human HCAR3, and the dashed red arrow depicts the overall orientation of superposed 3HO within the ligand-binding pocket of each HCAR2/3 model, as shown in A to G. Distinguishing residues between HCAR2 and HCAR3 are highlighted in blue.

**Supplementary Figure S8.**
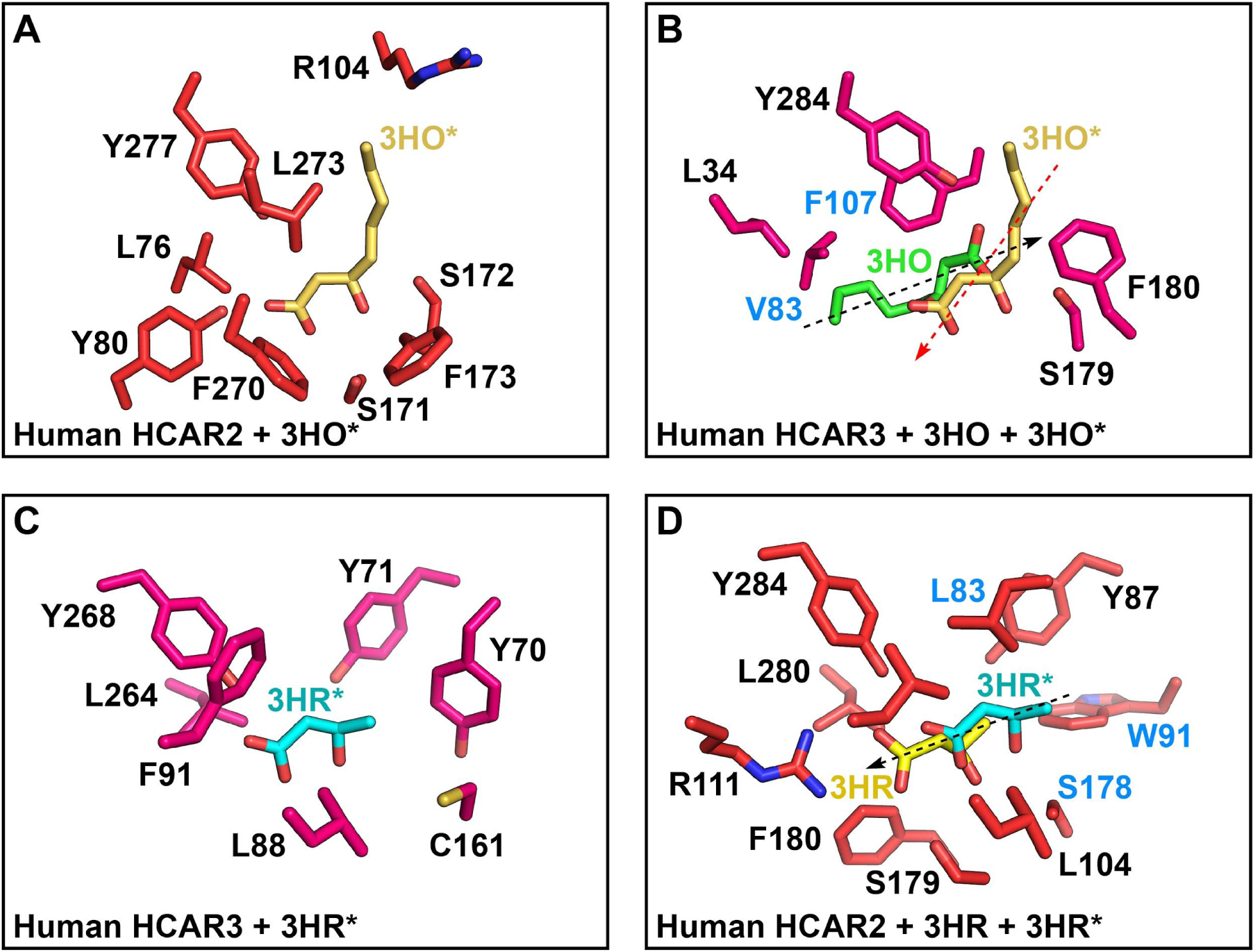
Comparison of ligand swapping in human HCAR2 and HCAR3. (A) Stick representation of the human HCAR2 ligand-binding pocket, with amino acid residues shown in red, bound to the non-canonical ligand 3-hydroxyoctanoate (3HO*; carbon atoms in yellow), as predicted by AlphaFold3. (B) Stick representation showing the superposed 3HO* ligand from panel A within the ligand-binding pocket of human HCAR3 (residues shown in hot pink). The spatial orientation of the canonical ligand 3HO (green carbons) is compared with the superposed 3HO* (yellow carbons). The dashed black arrow indicates the orientation of 3HO, whereas the dashed red arrow indicates the opposite axial orientation adopted by 3HO*. (C) Stick representation of the human HCAR3 ligand-binding pocket (hot pink carbons) bound to the non-canonical ligand β-hydroxybutyrate (3HR*; cyan carbons), as predicted by AlphaFold3. (D) Stick representation showing the superposed 3HR* ligand from panel C within the ligand-binding pocket of human HCAR2 (residues shown in red). The spatial orientation of canonical 3HR (yellow carbons) is compared with that of superposed 3HR* (cyan carbons). The dashed black arrow depicts the parallel axial orientation of 3HR and 3HR*.

### Divergent Expression Patterns of HCAR Paralogs are Consistent with Subfunctionalization

After gaining insight into the evolutionary and structural features of the HCAR gene family, we next investigated the evolutionary fate associated with their gene expression patterns in humans. To this end, we analyzed transcript abundance using data from the GTEx database (GTEx Consortium 2020), which revealed distinct tissue-specific expression profiles (Fig. 10). HCAR1 is predominantly expressed in breast, spleen, and adipose tissues, whereas HCAR2 and HCAR3 are highly expressed in whole blood and, to a lesser extent, in skin and esophageal tissues, with HCAR2 showing stronger expression than HCAR3 in the latter two tissues (Fig. 10A). Notably, HCAR1 expression is absent in the tissues where HCAR2 and HCAR3 are most highly expressed (Fig. 10A), suggesting a pattern of subfunctionalization. Within the hematopoietic cells, CD14⁺ monocytes were identified as the primary blood cells expressing HCAR2 and HCAR3, as determined by Cap Analysis of Gene Expression (CAGE) data (not shown). Given these differential expression patterns, we next examined the genomic context of the HCAR locus by integrating chromatin accessibility (ATAC-seq) and chromatin immunoprecipitation (ChIP-seq) datasets. ATAC-seq profiles from CD14⁺ monocytes and breast epithelium revealed that promoter accessibility mirrors gene expression (Fig. 10B). The HCAR1 promoter is open in breast tissue but closed in monocytes, whereas HCAR2 and HCAR3 promoters are accessible in monocytes, but not in breast tissue (Fig. 10B, blue shading). The CCCTC-binding factor (CTCF), a key architectural protein that regulates enhancer–promoter communication and delineates genomic domain boundaries (Cuddapah et al. 2009), was then analyzed to explore chromatin organization across the HCAR locus. By integrating CTCF ChIP-seq peaks with regions of accessible chromatin defined by ATAC-seq (Wernig-Zorc et al. 2024), we identified CTCF-binding sites flanking the HCAR cluster (Fig. 10B, pink shading). This arrangement indicates that the HCAR genes reside within a topologically associating domain (TAD) of approximately 100 kb, whose chromatin architecture and accessibility are differentially modulated in a tissue-specific manner (Fig. 10C).

**Figure 10.**
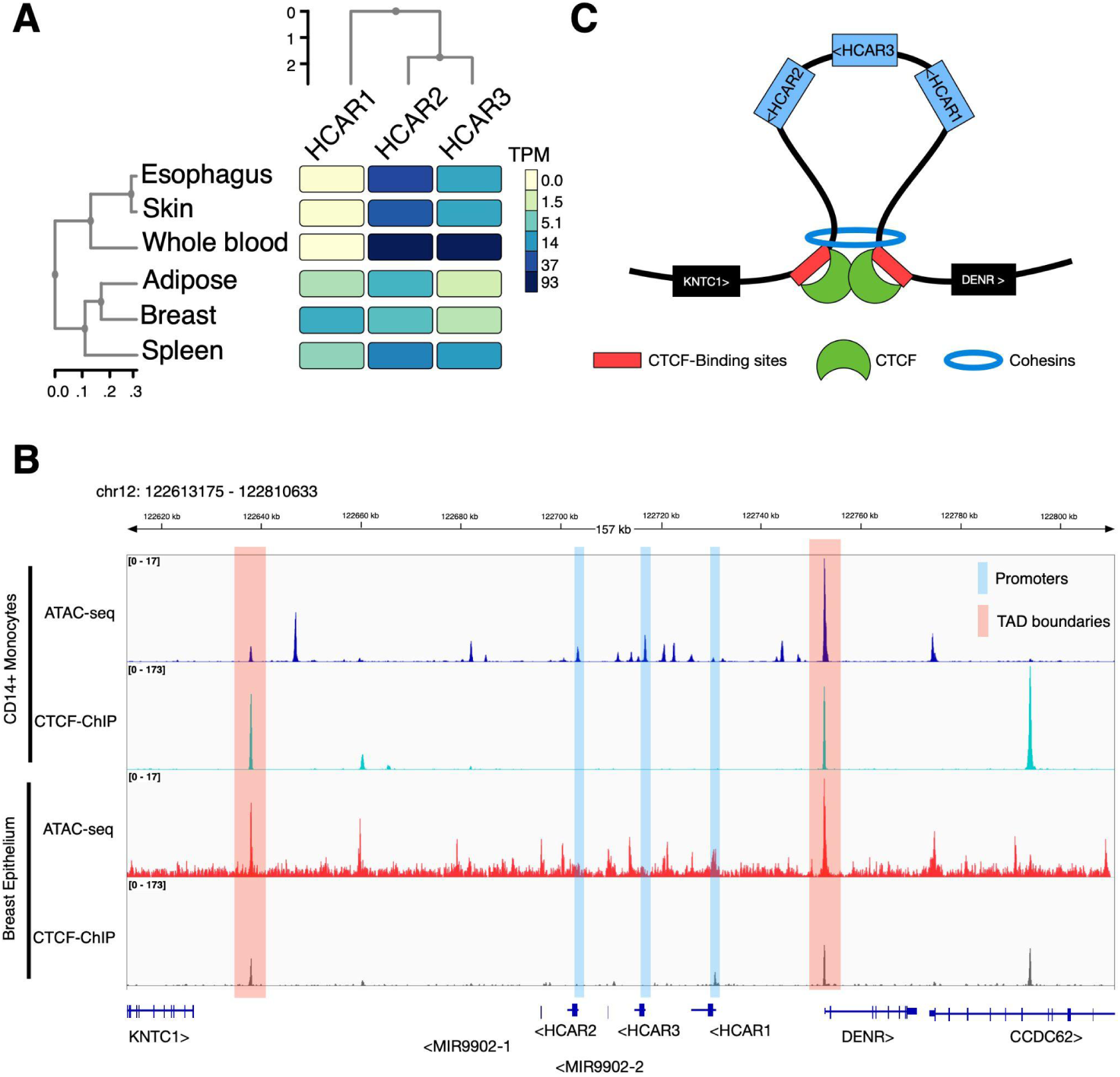
Expression patterns and chromatin organization of the human HCAR gene cluster. (A) Heatmap showing the expression levels (TPM, Transcripts Per Million) of HCAR1, HCAR2, and HCAR3 across selected human tissues (esophagus, skin, whole blood, adipose tissue, breast, and spleen). Hierarchical clustering reflects similarities in both tissue expression profiles and gene expression patterns. (B) Genomic view of the HCAR locus on chromosome 12. ATAC-seq and CTCF ChIP-seq profiles from CD14⁺ monocytes (accession numbers GSE254979 and ENCFF451AYU, respectively) and breast epithelium (accession numbers GSM5259179, and ENCFF378EET, respectively) are shown. Promoter regions are highlighted in blue and topologically associating domain (TAD) boundaries in pink, illustrating conserved regulatory features and chromatin accessibility across cell types. (C) Schematic representation of the three-dimensional chromatin architecture of the HCAR gene cluster, illustrating CTCF-binding sites (red), CTCF proteins (green), and cohesin complexes (blue). The loop structure places HCAR2, HCAR3, and HCAR1 within a shared regulatory domain flanked by KNTC1 and DENR genes.

Subfunctionalization through regulatory divergence, whereby paralogs become expressed in distinct tissues, is widely recognized as a common outcome of gene duplication (Ohno 1970; Lynch and Conery 2000; Lynch and Force 2000). A classic example is provided by the expression patterns of lactate dehydrogenase (LDH) isoforms. LDH is a tetrameric enzyme composed of subunits encoded by distinct paralogs, with LDHA (muscle type) and LDHB (heart type) representing the main somatic genes, and LDHC showing testis-specific expression (Claps et al. 2022; Farhana and Lappin 2023). Although LDH activity is present in all tissues, different combinations of these subunits give rise to tissue-specific isozymes. For example, the isoenzyme composed of four heart-type subunits (H₄) predominates in the heart, red blood cells, and germ cells, whereas the isoenzyme composed of four muscle-type subunits (A₄) is enriched in liver, skeletal muscle, and kidney (Claps et al. 2022; Farhana and Lappin 2023). Similar patterns of tissue partitioning are observed in other metabolic and transport gene families, including glucose transporters (SLC2A/GLUT) (Medina and Owen 2002; Mueckler and Thorens 2013), phosphofructokinase-1 (PFK) (Dunaway 1983), and aquaporins (AQP) (King et al. 2004; Finn and Cerdà 2015). Large-scale transcriptomic analyses, particularly from the GTEx consortium, demonstrate that such tissue-restricted expression of paralogs is widespread in the human genome and represents a common evolutionary outcome of gene duplication driven primarily by regulatory divergence rather than changes in protein function (Kryuchkova-Mostacci and Robinson-Rechavi 2016; GTEx Consortium 2020).

## Conclusion

We reconstructed the evolutionary history of the hydroxycarboxylic acid receptor gene family in primates, with a particular focus on the recently originated HCAR2 and HCAR3 gene lineages. Our phylogenetic analyses reveal that the duplicative history of these genes is more complex than previously proposed, involving multiple independent duplication events during ape evolution. We also show that non-ape primates retain the ancestral single-copy gene (HCAR2/3), whose structural, sequence, and ligand-binding analyses consistently indicate that it is likely to be functionally equivalent to HCAR2. Furthermore, expression and chromatin accessibility analyses demonstrate that major regulatory divergence occurred after the earlier split between HCAR1 and the HCAR2/3 gene lineages, while HCAR2 and HCAR3 retain largely overlapping expression profiles. Our results clarify homologous relationships within the HCAR gene family in primates, highlighting the importance of accurately reconstructing gene duplication histories for functional interpretation.

We demonstrate that HCAR3 originated independently in different ape lineages. The physiological meaning of this evolutionary innovation is beyond the scope of the present study, but some metabolic considerations may be warranted. HCAR2 and HCAR3 share intracellular signaling and tissue expression patterns, so novelty seems to lie elsewhere. The emergence of HCAR3 endowed cells with the ability to sense 3-hydroxyoctanoic, to the expense of β-hydroxybutyrate. Adipose tissue is made of triglycerides, which are oxidized to CO_2_ for the production of ATP, a complex branched pathway involving multiple tissues. Both 3-hydroxyoctanoic and β-hydroxybutyrate belong to this pathway, and both rise in plasma during fat mobilization. There are however important distinctions regarding information and how it is conveyed. Situated much higher in the catabolic chain, 3-hydroxyoctanoic reflects more directly the metabolic status of peripheral tissues (e.g. adipose tissue and muscle) than β-hydroxybutyrate, which is made later on in the liver, subject to strong factors like glucose and insulin. Another key difference is concentration. Circulating 3-hydroxyoctanoic levels are 100 times lower than those of the ketone body, resulting in shorter turnover time and better signal-to-noise ratio (Barros et al. 2013). Thirdly, β-hydroxybutyrate is not just a signal but an energy substrate, accounting for up to 30% of the budget in the starving brain (Achanta and Rae 2017), which is not the case for 3-hydroxyoctanoic. These differential features suggest that 3-hydroxyoctanoic is a more specialized signal of lipid mobilization, resembling the contrast between professional neurotransmitters like noradrenaline and serotonin and non-professional signals like lactate, β-hydroxybutyrate and adenosine (Barros et al. 2025).

For these reasons, species that possess HCAR3 have access to a more refined control of lipid metabolism compared with species that rely exclusively on HCAR2/3, like all non-ape species (Offermanns 2014; Offermanns 2017; Suzuki et al. 2023). While HCAR2 primarily senses β-hydroxybutyrate and mediates a broad antilipolytic response during prolonged fasting or deep ketosis, HCAR3 responds preferentially to intermediate lipid-turnover states (Offermanns 2014; Offermanns 2017; Suzuki et al. 2023). This additional receptor increases the resolution of Gi-mediated inhibition of lipolysis, providing a negative-feedback mechanism that prevents excessive fatty-acid release and energy waste when lipid availability is high but not ketogenic (Offermanns 2014; Offermanns 2017). In apes, this metabolic sensing refinement is particularly relevant given their distinctive energetic phenotype (Simmen et al. 2017). Compared with other primates, such as Old World monkeys, which lack HCAR3 and depend solely on HCAR2/3, hominids exhibit diets that are more energy-dense, lower in bulk, and more stable across seasons, together with tighter regulation of lipid mobilization to support large brains and prolonged life-history stages (Simmen et al. 2017). Thus, HCAR3 does not replace HCAR2 but complements it, endowing hominids with a more granular and robust lipid-sensing system aligned with their unique dietary strategies and metabolic demands. Finally, it has also been proposed that the main evolutionary advantage of retaining HCAR3 is that this receptor provides a novel system for sensing and mounting physiological responses to metabolites derived from lactic acid bacteria, particularly those originating from fermented foods and/or the gut microbiota (Peters et al. 2019). In this context, HCAR3 would function as an additional “sensor” linking diet-associated microbial metabolites to host physiology, with a particularly important role in shaping immune responses (Peters et al. 2019).

## Author Contributions

All authors contributed to the work presented in this paper. JCO, GAM, RM and KZ analyzed data. JCO, GAM, RM, and LFB wrote the paper.

## Acknowledgments

This work was supported by Fondo Nacional de Desarrollo Científico y Tecnológico from Chile, FONDECYT 1250688 to JCO, FONDECYT 1230145 to LFB, FONDECYT 1251655 to GAM, and Fondo de Fomento al Desarrollo Científico y Tecnológico, FONDEF ID25I10174 to RM.

## Data Availability

Data and supplementary material are available online at https://github.com/opazolab/HCAR

## Supplementary Material

Supplementary material is available online.

